# Evidence that the TRPV1 S1-S4 Membrane Domain Contributes to Thermosensing

**DOI:** 10.1101/711499

**Authors:** Minjoo Kim, Nicholas J. Sisco, Jacob K. Hilton, Camila M. Montano, Manuel A. Castro, Brian R. Cherry, Marcia Levitus, Wade D. Van Horn

**Author notes:** These Authors contributed equally to this work (Minjoo Kim, Nicholas J. Sisco).

## Abstract

Sensing and responding to temperature is crucial in biology. The TRPV1 ion channel is a well-studied heat-sensing receptor that is also activated by vanilloid compounds including capsaicin. Despite significant interest, the molecular underpinnings of thermosensing have remained elusive. The TRPV1 S1-S4 membrane domain couples chemical ligand binding to the pore domain during channel gating. However, the role of the S1-S4 domain in thermosensing is unclear. Evaluation of the isolated human TRPV1 S1-S4 domain by solution NMR, Far-UV CD, and intrinsic fluorescence shows that this domain undergoes a non-denaturing temperature-dependent transition with a high thermosensitivity. Further NMR characterization of the temperature-dependent conformational changes suggests the contribution of the S1-S4 domain to thermosensing shares features with known coupling mechanisms between this domain with ligand and pH activation. Taken together, this study shows that the TRPV1 S1-S4 domain contributes to TRPV1 temperature-dependent activation.

## Introduction

Transient receptor potential (TRP) ion channels are a family of membrane proteins which play diverse roles in physiology (*1, 2*). TRPV1, from the vanilloid subfamily, is responsive to various chemical and physical stimuli, including vanilloid ligands, elevated temperature, protons, endogenous lipids, and small modulatory membrane proteins (*3–6*). While TRPV1 is expressed in neuronal and non-neuronal tissues, its role in neuronal tissues has garnered significant interest (*7*). For example, in group C unmyelinated nerve tissue of the peripheral nervous system, TRPV1 is integral to nociception (pain) (*8, 9*). Consequentially, there is interest in TRPV1 therapeutic intervention for various pain indications (*8*). Beyond nociception, an increasing number of studies provide emerging evidence that TRPV1 is involved in diverse human physiologies and pathophysiologies including: inflammatory diseases (*10–12*), obesity (*13, 14*), diabetes (*11*), longevity (*15*), and cancer (*16, 17*).

Cryo-electron microscopy (cryo-EM)-based structural biology has had significant impact on understanding the molecular architecture that underlies TRP channel function (*18*). These structures have shown that TRP channels resemble the transmembrane topology of voltage-gated ion channels (VGICs), with two conserved transmembrane structural domains. The S1-S4 transmembrane helices (S1-S4 domain) form a four-helix bundle that is structurally related to the voltage-sensing domain (VSD) in VGICs. In TRP channels, like their evolutionary ancestors in the VGIC superfamily, the S5-S6 transmembrane helices form the pore domain (PD), which assembles into a tetrameric channel. In addition to the identifying topological features, a number of TRPV1 structures in different states have been determined, which have provided significant insight into the molecular basis for TRPV1 chemical activation (*19–21*). Canonical TRPV1 vanilloid compounds, like the pungent agonist capsaicin, bind to the S1-S4 domain which couples with the PD to open the lower and upper gates of the channel (*22*), thereby initiating signal transduction. The cryo-EM determined vanilloid binding site (*19–21*) is consistent with previous studies (*23–28*), which identified residues in the S3 and S4 helices of the TRPV1 S1-S4 domain as central for capsaicin activation.

Several TRP channels are exquisitely sensitive to changes in temperature and function as molecular thermometers. The temperature-induced activation of thermosensitive TRP channels generates large changes in enthalpy (*ΔH*) and significant compensating changes in entropy (*ΔS*), resulting in biologically accessible changes in free energy (*ΔG*) between closed and open states. Typically to assess thermosensitivity of biological systems, the temperature coefficient (Q_10_) is measured, which is simply the ratio of equilibrium constants measured 10 °C apart. The temperature coefficient is related to the change in enthalpy (*29*), and thermosensitive TRP channels like TRPV1 have large Q_10_ and *ΔH* values of ~40 and ~100 kcal/mol, respectively (*30, 31*). The physical mechanisms that underlie temperature sensitivity are thought to arise from changes in secondary structure and hydrophobic/hydrophilic accessibility, both of which have large thermodynamic signatures (*1, 29, 32, 33*). Notwithstanding, the region (or regions) within TRP channels that are key to thermosensing remains controversial. As a result, the specific biochemical and structural mechanisms that give rise to TRP channel temperature sensing are also not well understood. However, important contributions have been made to investigating TRPV1 thermosensing. For example, studies of the purified TRPV1 channel clearly show that the channel is intrinsically thermosensitive (*34*). Similarly, an isoform of murine TRPV1 that lacks the majority of the N-terminus remains thermosensitive (*35*), suggestive that the transmembrane region is central to thermosensing. More recently, a chimeric study which swapped the TRPV1 PD into a non-thermosensing channel, was able to endow thermosensitivity in the chimeric channel (*36*). Despite these studies, the mechanism of TRPV1 thermosensing has remained elusive.

Given the role of the TRPV1 S1-S4 domain (V1-S1S4) in ligand activation and the structural similarity to voltage-sensing domains in VGICs, we hypothesize that the S1-S4 domain may contribute to the thermosensitivity of TRPV1. We adopt a distinct strategy that is not reliant on mutagenesis but instead focuses on direct temperature-dependent characterization of the hV1-S1S4, an evolutionarily conserved structural domain (Fig. 1). Here, we show that an isolated hV1-S1S4 is in a biologically relevant state, is sufficient for vanilloid ligand binding, and retains the expected secondary structure and membrane topology. Temperature-dependent studies of the hV1-S1S4 using solution nuclear magnetic resonance (NMR) spectroscopy identify a two-state transition between folded conformational states. The magnitude of the NMR detected temperature sensitivity is supported by far-UV circular dichroism (CD) and intrinsic tryptophan fluorescence spectroscopy. Quantitative comparison with whole-cell patch-clamp electrophysiology experiments in mammalian cells indicates that the hV1-S1S4 significantly contributes to TRPV1 thermosensitivity. Additionally, the temperature-dependent conformational changes were examined by NMR-based temperature-dependent distance measurements from paramagnetic relaxation enhancement (PRE), secondary structure measurements from chemical shift assignment and residual dipolar coupling, and solvent exposure from deuterium/hydrogen exchange and water–protein NOEs. To validate these outcomes, we introduced a mutation that disrupts coupling between the S1-S4 and pore domains and show that both ligand and temperature activation are abrogated in the full-length channel but retained in the isolated domain. Taken together, the data provide thermodynamic and mechanistic insight into the properties of thermosensitive TRP channels and suggests an overlap between TRPV1 ligand, proton (pH), and temperature activation.

**Figure 1.**
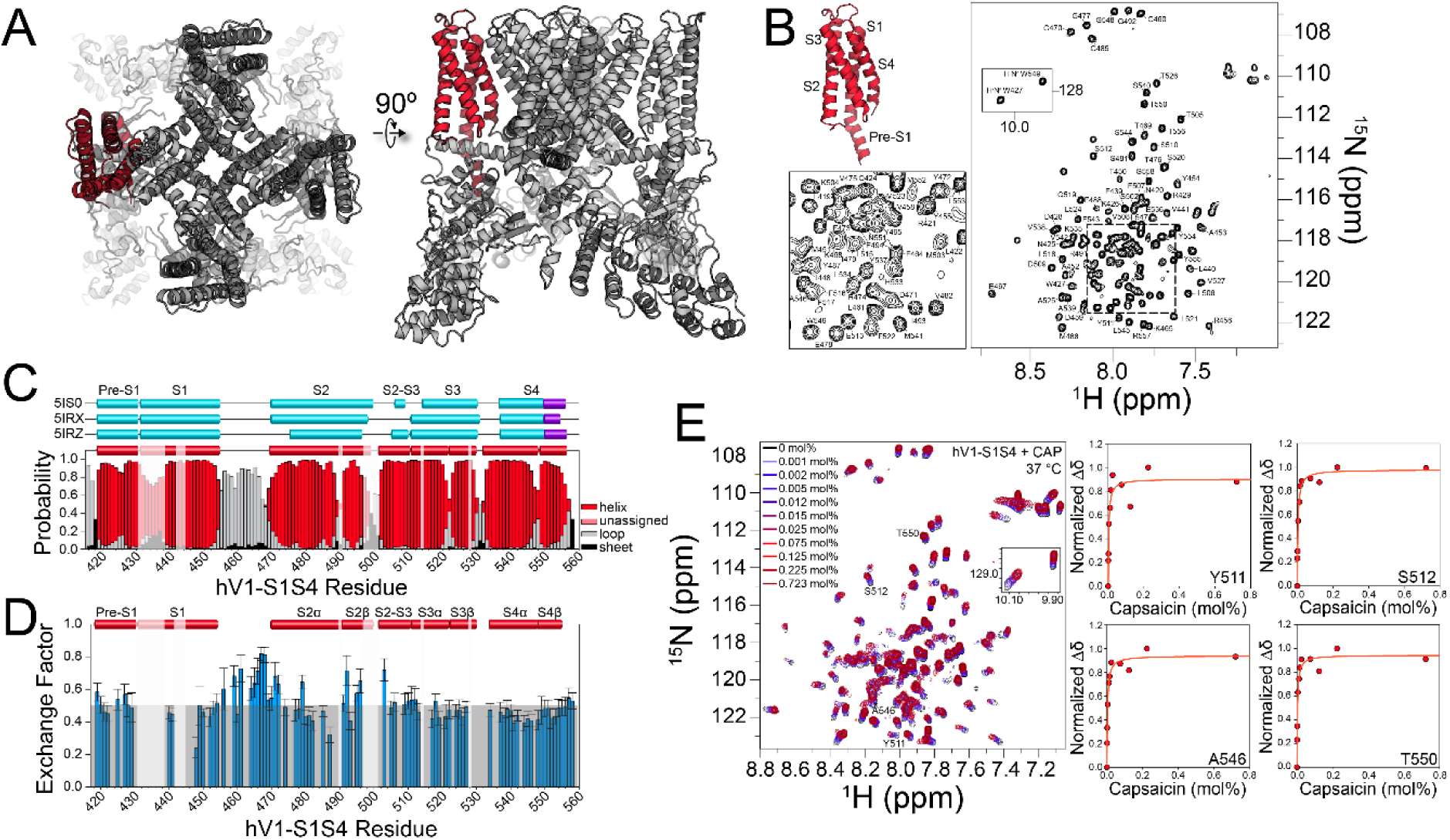
The isolated human TRPV1 S1-S4 domain is folded in a biologically relevant state and binds capsaicin at elevated temperatures. (**A**) Highlighted in red is the hV1-S1S4 construct shown in the rTRPV1 cryo-EM structure (PDB ID: 3J5P) and used throughout our studies. (**B**) Representative ^15^N-TROSY-HSQC showing the hV1-S1S4 backbone resonance assignments at 45 °C. (**C**) Cryo-EM determined secondary structure information of rTRPV1 (cyan/purple) compared with the NMR determined secondary structure (red) from hV1-S1S4 at 45 °C identifies the similarities and differences in secondary structure. The S4 3_10_ helix is shown in purple. (**D**) Deuterium/hydrogen (D/H) exchange factors show the hV1-S1S4 solvent accessibility is consistent with the anticipated membrane topology. (**E**) Superimposed ^1^H-^15^N HSQC spectra of the hV1-S1S4 titration with capsaicin at 37 °C. Canonical capsaicin binding residues include Y511, S512, and T550, and A546, show saturating binding isotherms as a function of capsaicin.

## Results

### The isolated hV1-S1S4 resides in a biologically relevant state

#### The hV1-S1S4 in isolation retains the expected membrane topology

The transmembrane hV1-S1S4 construct includes the pre-S1 helix through the S4 helix (residues 417 to 558) (Fig. 1A). Optimization of hV1-S1S4 expression resulted in ~1.5 mg of purified protein per liter of M9 media. The identity of the hV1-S1S4 was verified by SDS-PAGE, western blot, LC-MS/MS, and ultimately NMR resonance assignment (fig. S1, A to C). After membrane mimic screening, the hV1-S1S4 was reconstituted in LPPG (1-palmitoyl-2-hydroxy-*sn*-glycero-3-phospho-(1’-*rac*-glycerol)) lysolipid micelles, which yielded high-quality NMR spectra (fig. S1D). We focused on single chain membrane mimics, such as LPPG, to avoid temperature-dependent phase transitions common to lipids that could interfere with temperature-dependent thermodynamic and biophysical characterization. TROSY-based 3D and 4D NMR experiments were carried out to assign the hV1-S1S4 amide backbone. Specifically, TROSY-based HNCA, HNCOCA, HNCO, HNCACB, CBCACONH, HNCACO, ^15^N-edited-NOESY-TROSY, 4D HNCACO and 4D HNCOCA were used to assign 87% of backbone resonances (BMRB ID: 27029, Fig. 1B and fig S1, E to J). The hV1-S1S4 membrane topology was determined from secondary structure derived from backbone chemical shift data (TALOS-N) and membrane accessibility from NMR-detected deuterium-hydrogen (D/H) exchange factors (Fig. 1, C and D). The resulting hV1-S1S4 secondary structure in solution at elevated temperature is similar to existing rat TRPV1 cryo-EM structures. Similarly, the D/H exchange factors show that the transmembrane helices have lower exchange factors than the solvent accessible loop regions, indicative of the expected membrane topology for this domain.

#### The hV1-S1S4 domain specifically binds capsaicin

The isolated hV1-S1S4 solubilized in LPPG retains the ability to specifically bind capsaicin at 37 °C (Fig. 1E). Previously identified capsaicin binding residues, Y511, S512, and T550, were used as probes of capsaicin binding. These residues showed saturable chemical shift perturbation as a function of increased capsaicin concentration which is indicative of specific binding (Fig. 1E). The average *K*_d_ values from the key vanilloid binding residues (Y511, S512, and T550) is 3.4 ± 0.4 mmol%. L547 (M547 in the rat ortholog) has also been ascribed importance in vanilloid binding and activation; however, due to resonance overlap this residue could not be analyzed. A neighboring residue, A546, which is located within 5 Å of vanilloid ligand as shown a vanilloid-bound TRPV1 structure (PDB ID: 5IRX), also exhibited saturating binding with a *K*_d_ value of 3.6 ± 0.5 mmol% (Fig. 1E). Taken together, the isolated hV1-S1S4 in solution retains four transmembrane embedded helices and recapitulates expected features of capsaicin binding, indicating that the isolated hTRPV1-S1S4 retains a biologically-relevant conformation in solution and at elevated temperatures.

### Two-state temperature transition and thermodynamic analysis of the full-length human TRPV1

TRPV1 temperature-dependent electrophysiology data have been shown to fit well to a simple two-state thermodynamic model between open (active) and closed (resting) states (*2*). While this is certainly an oversimplification, modeling TRPV1 temperature-dependent transitions allows for quantification of the thermosensitivity. In this two-state model the temperature-dependent slope of the transition between conformational states reflects the change in enthalpy (Δ*H*) (*37*) and is a read out of the thermosensitivity associated with the conformational change between temperatures.

To date the majority of TRPV1 thermosensing studies have focused on rodent orthologs (Table S1). To evaluate the thermosensitivity of the human TRPV1 ortholog, whole-cell patch-clamp electrophysiology measurements were performed using the full-length wild-type human TRPV1 in HEK293 cells. We employed two distinct methods to measure the thermosensitivity (*ΔH*) of human TRPV1 in cellular conditions. First, steady-state current values were measured at +60 mV as a function of increasing temperature (fig. S2A). The normalized current values were plotted against temperature and fit to a two-state model representing closed (resting) and open (active) states resulting in a measured Δ*H* of 98 ± 12 kcal/mol (Fig. 2A). Independently, we used a second method involving temperature ramps from 20 to 50 °C and fit the data to a pseudo-steady state thermodynamic model to obtain hTRPV1 Δ*H* values (see Methods), resulting in an average Δ*H* of 94 ± 8 kcal/mol (fig. S2B). The Δ*H* values of human TRPV1 measured from the steady-state current and the temperature ramp methods are consistent with those of rodent TRPV1 from previous studies, which is ~90 kcal/mol (Table S1).

**Figure 2.**
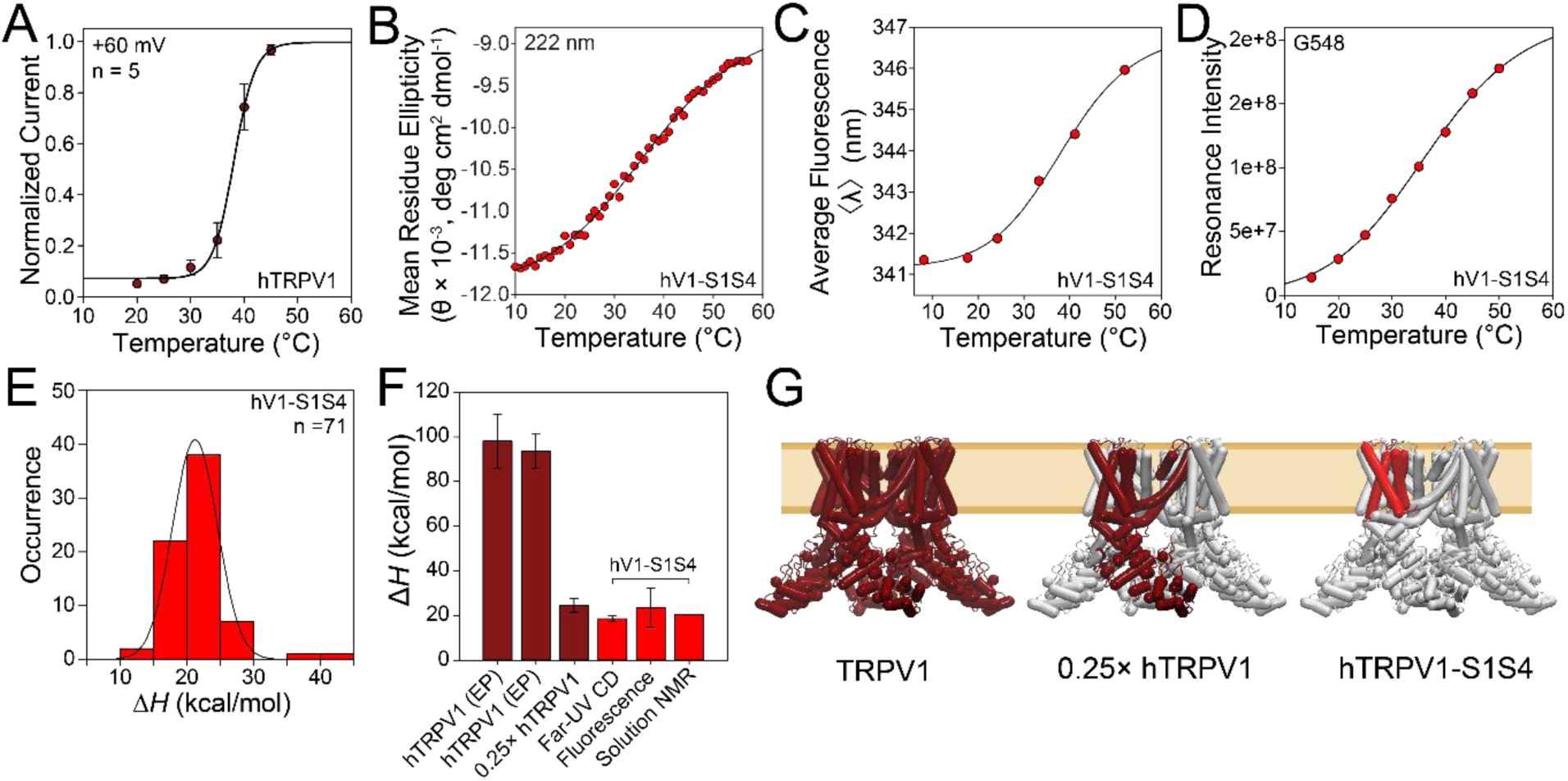
Temperature-dependent two-state behavior and thermodynamic analysis of hTRPV1 and hV1-S1S4. (**A**) Whole-cell patch-clamp electrophysiology measurements from a full-length human TRPV1 in HEK293 cells. (**B**) Temperature ramp of hV1-S1S4 monitored by far-UV circular dichroism shows two-state behavior. (**C**) The first moment of intrinsic tryptophan fluorescence of hV1-S1S4 as a function of temperature shows two-state transition similar to that identified by CD. (**D**) Representative NMR resonance intensity, G548 in the S4 helix, shows two-state behavior. G548 was chosen due to its similarity to the mean Δ*H* obtained from NMR. (**E**) A histogram of 71 enthalpies from NMR resonance intensities are shown in red. These data fit to a Gaussian function (black line) provide an average value of 21.2 ± 0.1 kcal/mol (**F**) Comparison of Δ*H* values, the first dark red two bars are from the cellular electrophysiology measurements for full-length hTRPV1 from steady state currents (**A**) and temperature ramps. A representative value for one quarter of the steady state value is also shown. The Δ*H* values from CD, fluorescence, and NMR are shown in red. (**G**) The left panel shows the full-length TRPV1 used for electrophysiology measurements. The middle panel shows the relative size of a TRPV1 monomer. The right panel shows the hV1-S1S4 domain in red, which has similar Δ*H* as the monomer, showing the relative significance of the hV1-S1S4 in thermosensitivity.

### Two-state temperature transition and thermodynamic analysis of the hV1-S1S4 domain

To investigate a potential role of the hV1-S1S4 in thermosensitivity, far-UV circular dichroism (CD) and solution NMR spectra were recorded at 20 °C and 50 °C; temperatures that would be expected to correspond to distinct conformational states if the hV1-S1S4 were to contribute to thermosensitivity. First, the CD spectra at 20 °C and 50 °C displayed characteristic α-helical minima at 208 and 222 nm. These spectral features were retained at both high and low temperatures, suggestive that the protein does not significantly unfold under these conditions (fig. S2, C and D). Additionally, the ^1^H-^15^N TROSY-HSQC NMR spectra of hV1-S1S4 at these two temperatures (20 °C and 50 °C) provide qualitative insight into potential temperature-dependent conformational change and the foldedness of this domain at the respective temperatures (fig. S2E). The NMR proton resonance dispersion and resolution indicate that the hV1-S1S4 remains generally folded over this temperature range. There is also significant temperature dependent chemical shift perturbation between the spectra at 20 °C and 50 °C, which is consistent with a temperature-induced conformational change. The temperature-dependent spectral changes, in both CD and NMR, are completely reversible and indicate that the system is suitable for thermodynamic analysis to assess thermosensitivity (Δ*H*) of the apparent conformational change (fig. 2, C to E).

#### CD-based temperature-dependent studies of hV1-S1S4

CD data collected as a function of temperature have long been used to determine thermodynamic properties of proteins (*37*). The hV1-S1S4 CD data exhibit two-state behavior at 222 nm, near the characteristic α-helical minimum, as a function of temperature. Fitting the data to a two-state sigmoidal model yielded a *ΔH* of 19 ± 1 kcal/mol, which reflects the temperature dependence of the hV1-S1S4 change conformational states (Fig. 2B). The KCNQ1-voltage-sensing domain (VSD, helices S1-S4) is structurally homologous to the hV1-S1S4; however, it is not directly activated (gated) by thermal stimulus. As a control, we measured the KCNQ1-VSD mean residue ellipticity (MRE) at 222 nm over the same temperature range. The hKCNQ1-VSD did not exhibit two-state behavior, but instead showed a linear temperature response over this temperature range, consistent with a general thermal expansion of the α-helical hydrogen bonds (fig. S3A). Comparatively, the lack of measurable thermosensitivity (Δ*H*) of the KCNQ1-VSD over this temperature range suggests that the hV1-S1S4 reflects a feature of TRPV1 thermosensitivity.

#### Intrinsic tryptophan fluorescence temperature-dependent studies of hV1-S1S4

The hV1-S1S4 was also subjected to temperature-dependent studies using intrinsic tryptophan fluorescence. Typically, intrinsic tryptophan has a large fluorescence transition dipole moment and the position and shape of the emission spectrum is particularly sensitive to the polarity of the environment. Tryptophan (Trp) emission maxima in an apolar (hydrophobic) environment is blue-shifted and varies from ca. 310-335 nm. Whereas emission maxima in polar (hydrogen-bonding) solvents for an unstructured Trp residue is red-shifted about 40 nm up to ca. 355 nm (*38*). These characteristics are commonly exploited in protein folding studies, where the emission peak of Trp shifts towards the red as the fraction of unfolded protein increases. In the context of hV1-S1S4, this analysis allows insight into both the thermosensitivity of the transition and the environment of the Trp residues. The hV1-S1S4 has two endogenous Trp residues in the amphipathic pre-S1 helix and the transmembrane S4 helix, both of which are anticipated to be in a hydrophobic environment from cryo-EM structures. Indeed the resulting fluoresce emission spectral maxima are 328 nm at 7.4 °C and 332 nm at 52.0 °C, indicating that over the temperature range studied, the endogenous Trp residues generally retain their membrane embeddedness (fig. S3). This observation is consistent with the observations from CD (fig. S2C) and NMR (fig. S2D) that the hV1-S1S4 remains generally folded across this temperature range. Quantifying the observed temperature induced spectral shifts in terms of the average emission wavelength (⟨λ⟩) identifies a sigmoidal two-state behavior with a Δ*H* of 24 ± 9 kcal/mol (Fig. 2C). This thermosensitivity is consistent with the observed CD-detected thermosensitivity (Fig. 2B).

#### NMR-detected temperature-dependent studies of hV1-S1S4

A series of ^1^H-^15^N TROSY-HSQC spectra of hV1-S1S4 were collected at temperatures ranging from 15 °C to 50 °C (fig. S3D). The hV1-S1S4 resonance assignments at 45 °C (Fig. 1B) were used to assign the temperature series spectra by chemical shift mapping, resulting in assignment of 71 resonances (~51% of the total expected resonances) across the temperature range (15 to 50 °C). The remaining resonances could not be unambiguously identified at all temperatures because of loss of resonance intensity, emergence of resonances, and/or coalescence with other resonances. The resonances that could be assigned across the temperature series show reversible two-state temperature-dependent behavior, as illustrated by representative plots of G548 (Fig. 2D) and W549 (fig. S3E). The two-state temperature-dependent behavior is similar to that observed from CD and fluorescence emission experiments. Fitting the 71 assigned resonances as a function of temperature to the same two-state model used for electrophysiology, CD, and fluorescence, enabled us to evaluate the Δ*H* value for each individual resonance. Plotting the Δ*H* values as a function of frequency (Fig. 2E) suggests the mean Δ*H* from NMR has similar magnitude as those observed from CD and fluorescence measurements. Fitting these data to a Gaussian distribution results in an average thermosensitivity of Δ*H* = 21.2 ± 0.1 kcal/mol and a mean T_50_ (reflective of the inflection point of the sigmoid) of 40.7 ± 0.6 °C (Fig. 2E and fig. S3F). To ensure that the effects observed by NMR, CD, and fluorescence arise from changes in temperature and not pH, the experiments were carried out in sodium phosphate buffer, which is relatively insensitive to temperature-induced pH changes. We measured the change in pH to be ~0.1 units when the temperature of the buffer is varied from 10 to 60 °C (Table S2). Also, hV1-S1S4 ^1^H-^15^N TROSY-HSQC NMR spectra show no detectible chemical shift perturbation over this pH range, pH 6.5 and 6.4 (fig. S3G). Taken together, the far-UV CD, intrinsic tryptophan fluorescence, and solution NMR data indicate that the hV1-S1S4 domain undergoes a temperature-dependent conformational change with a thermosensitivity of ~20 kcal/mol.

### Insights into temperature-dependent conformational change

#### Differences in secondary structure between rTRPV1 and hV1-S1S4

The secondary structure of the hV1-S1S4 at 45 °C was determined from experimental backbone chemical shift data analyzed in TALOS-N (Fig. 1C) (*39*). Comparison of secondary structures of the isolated hV1-S1S4 in solution at elevated temperature with those of the truncated rat TRPV1 structures from cryo-EM (*19–21*) indicate that the transmembrane topologies are generally consistent (Fig. 1C). Despite the similarities, there are differences in helicity between the NMR-determined secondary structure of hV1-S1S4 at 45 °C and the low temperature cryo-EM structures (Fig. 3A). The NMR data indicate that the S2, S3, and S4 helices have short helical kinks that are absent from the TRPV1 cryo-EM structures. Among the cryo-EM structures there are also differences; for example, in the apo structure (PDB ID: 5IRZ), which is the putative resting state structure, the S2 helix begins near R474. However, in the putative ligand-bound structures (PDB ID: 5IS0 and 5IRX) the extracellular side of the S2 helix is about two helical turns longer and begins near residue K468. The NMR data at elevated temperature indicates that the S2 helical conformation reflects that of the ligand bound structures. The NMR data also identify distinct helical content in the S2-S3 linker region, which is at the heart of the vanilloid binding pocket. In the apo and antagonist-bound cryo-EM structures (PDB IDs: 5IRZ and 5IS0 respectivly), the S2-S3 linker adopts a short amphipathic helix that includes residues S505-S510 with a break prior to the start of the S3 helix. Whereas, in the agonist activated structure (PDB ID: 5IRZ), the S2-S3 linker is not helical. The NMR-determined secondary structure is distinct and shows a near-continuous helix that begins near S502 in the S2-S3 linker and continues through the S3 helix. Additionally, the rTRPV1 structures show that the intracellular side of the S4 helix adopts a 3_10_ helical conformation (Fig. 1C). In solution at elevated temperatures, the bottom of S4 helix does not adopt a 3_10_ helix but instead has decreased helical probability, most similar to the agonist bound S4 helix (Fig. 1C).

**Figure 3.**
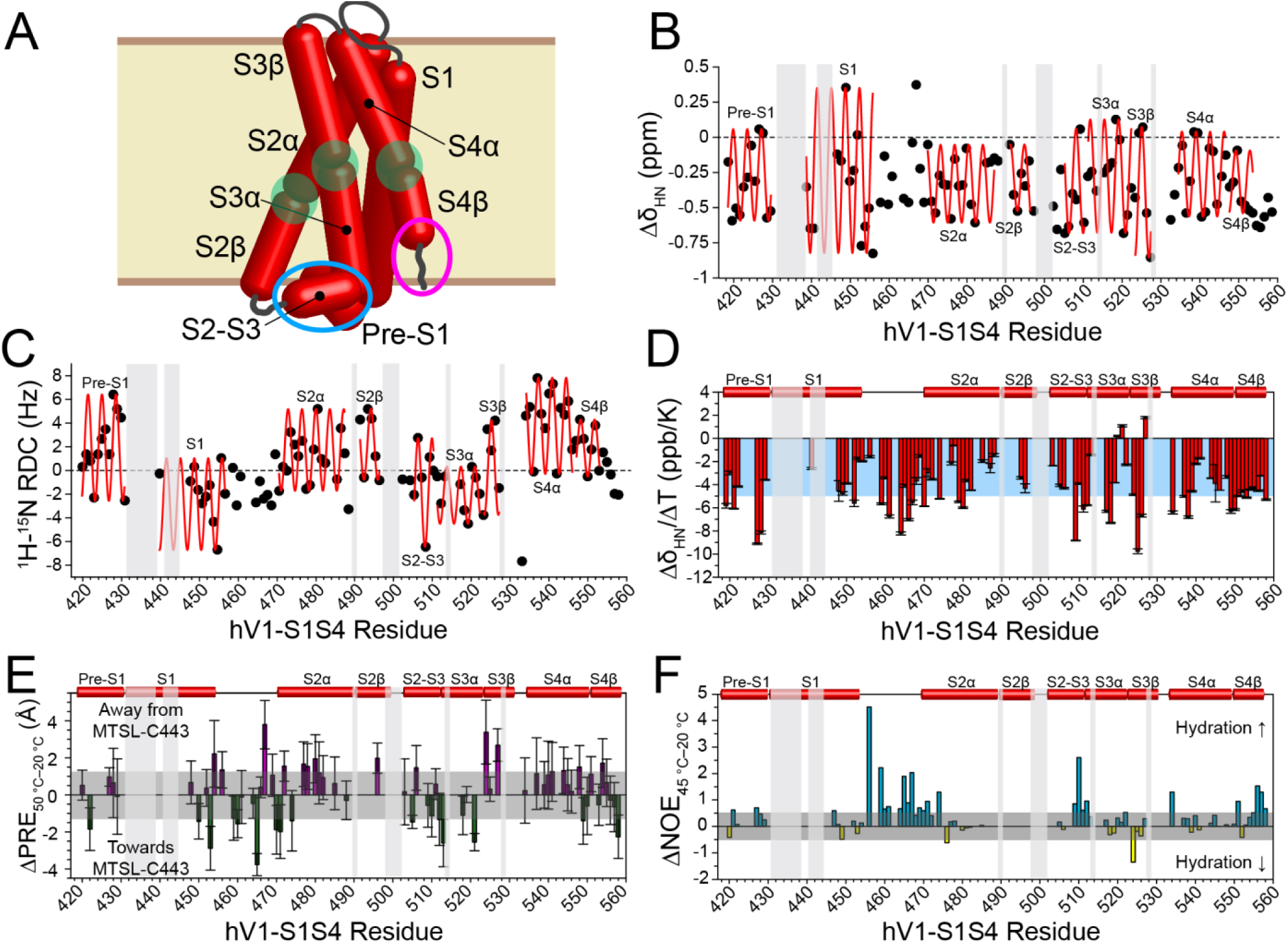
hV1-S1S4 undergoes subtle temperature-induced changes in secondary structure, distances, and solvent accessibility. (**A**) A cartoon of hV1-S1S4 secondary structure based on the TALOS-N prediction highlights secondary structural features that are distinct from cryo-EM rTRPV1 structures. Blue and magenta circles indicate increased and decreased helicity, respectively. Breaks in transmembrane helices are shown as transparent green circles. (**B**) Proton Δδ analysis identifies hV1-S1S4 secondary structure at elevated temperature (45 °C). Light grey boxes indicate that these regions where NMR resonances are unassigned. (**C**) Dipolar wave analysis from RDC measurements of the hV1-S1S4 at 45 °C identifies secondary structure. (**D**) A bar plot of the averaged amide proton temperature coefficients (Δδ_HN_/ΔT) against the hV1-S1S4 residue number. Values greater than −4.6 ppb/K are indicative of hydrogen bonding and are highlighted (blue). (**E**) Temperature dependent differences in distances measured using paramagnetic relaxation enhancement (ΔPRE) of the hV1-S1S4 at 20 °C and 50 °C. The average of the SEM, 1.2 Å, was used as a threshold value and shown in grey. (**F**) Changes in hydration as a function of temperature were measured from normalized H_N_–water resonance intensities from ^15^N-NOESY-TROSY data at 20 °C and 45 °C. The ΔNOE_45 °C−20 °C_ is reflective of changes in hydration where loops and the bottom of S4 helix exhibits increased hydration at elevated temperatures.

To further investigate and validate the apparent differences in helicity, we analyzed the secondary amide proton chemical shifts (Δδ_HN_, Fig. 3B). The proton Δδ values were calculated by taking the difference between an assigned resonance ^1^H chemical shift and the same amino acid in a random coil conformation. Δδ_HN_ data typically show a characteristic periodicity in helical regions (*40–42*). Changes in Δδ_HN_ were plotted against the hV1-S1S4 residue numbers, and each helix can be fit to a sinusoidal function for α-helices comprising the following amino acids: 419 to 430 for pre-S1, 439 to 456 for S1, 470 to 487 for S2α, 490 to 496 for S2β, 505 to 510 for S2-S3 amphipathic loop, 511 to 522 for S3α, 523 to 527 for S3β, 534 to 546 for S4α, and 547 to 553 for S4β. These data for hV1-S1S4 are consistent with the secondary structure output from TALOS-N. Additionally, the Δδ_HN_ data suggest that the intracellular side of the S4 helix, including residues L553 to G558, is not helical at higher temperature. This is distinct from the consensus 3_10_ helix identified over this region in the cryo-EM rTRPV1 structures. The RMSE values for corresponding helices are 0.26, 0.49, 0.18, 0.15, 0.33, 0.27, 0.22, 0.12, and 0.19 (Fig. 3B). Helicity can also be determined from NMR residual dipolar coupling (RDC) measurements, where fitting of the data to a helical sinusoid model can be used to identify alpha-, 3_10_-, and pi helices. This dipolar wave analysis of RDC data can also differentiate from ideal and kinked helices (*43*). ^1^H-^15^N RDCs were measured for hV1-S1S4 at 45 °C and used in dipolar wave analysis (Fig. 3C and fig. S4A). The same helices used in the proton Δδ analysis were fit to the same sinusoidal function and the RMSE values (Hz) for these fittings are 2.85, 2.51, 2.08, 1.18, 1.06, 1.31, 2.84, 2.43, 0.69, and 1.42. The outcomes of the RDC data further validate the TALOS-N and Δδ_HN_ chemical shift-based secondary structure identified for the isolated hV1-S1S4 domain in solution at elevated temperatures.

In RDC data, helical distortions can be identified by changes in dipolar wave amplitudes, which are a manifestation of alignment tensor magnitude. Using this metric, the RDC data indicate that hV1-S1S4 in solution has helical distortions in S2, S3, and S4 transmembrane helices consistent with the chemical shift data (Fig. 3C). RDC analysis of the intracellular S4 helix confirms a lack of helicity at elevated temperature consistent with the Δδ_HN_ data. We note that this region of the S4 segment in both the RDC and Δδ_HN_ data, beginning from L553, do not fit well to any helical subtype, including α-, 3_10_-, and π helices suggesting that this region in solution at elevated temperatures is distinct from that in the rTRPV1 cryo-EM structures.

Amide proton chemical shifts (δ_HN_) have a characteristic temperature dependence. The relationship between δ_HN_ and temperature is fit to a linear function where the slope of the fit is called the temperature coefficient (Δδ_HN_/ΔT). Δδ_HN_/ΔT values are an indicator of the probability of hydrogen bond formation and generally reflect secondary structure in helical membrane proteins (*44, 45*). More specifically, residues that form intramolecular hydrogen bonds typically have Δδ_HN_/ΔT values that are higher than −4.6 ppb/K (*44*). The temperature coefficients of the hV1-S1S4 were calculated and plotted against the residue number (Fig. 3D). Transmembrane helices had an average Δδ_HN_/ΔT value of −3.8 ± 0.3 ppb/K, indicating a more structured, hydrogen-bonded state; whereas, the extracellular/intracellular loops had a more negative average Δδ_HN_/ΔT value of −5.2 ± 0.5 ppb/K which is indicative of a lack of hydrogen bonding (Fig. 3D). Coupling the secondary structure analysis of the hV1-S1S4 domain from TALOS-N, Δδ_HN_, and ^1^H-^15^N RDCs, with Δδ_HN_/ΔT and D/H exchange factor data (Fig. 1, C and D, and Fig. 3, B to D) shows that in solution at elevated temperatures this domain retains the expected transmembrane topology, but with particular changes in helicity that either generally reflect the agonist activated cryo-EM structure or are distinct from the rTRPV1 cryo-EM structures.

#### Temperature-dependent distance changes in hV1-S1S4

To probe the magnitudes and directions of conformational change detected in the temperature titration studies of hV1-S1S4, NMR-detected paramagnetic relaxation enhancement (PRE) distance measurements were made from proton relaxation experiments at 20 °C and 50 °C. These experiments exploit the lone endogenous cysteine residue (C443, S1 helix) in hV1-S1S4 for MTSL labeling, obviating the need to engineer mutations that might impact the potential temperature-dependent structural changes. To ensure the highest possible accuracy of these measurements, six separate samples were prepared, the paramagnetic (three samples) and diamagnetic species (three samples) were each recorded in triplicate at each temperature. To accurately convert the PRE data to distance information, the rotational correlation times (τ_c_) were also directly measured at both temperatures using TRACT (<u>T</u>ROSY for rot<u>a</u>tional <u>c</u>orrelation times) NMR experiments (*46*). Using exclusively the W549 indole amine resonance, which is membrane embedded in the middle of the S4 transmembrane helix, the hV1-S1S4 rotational correlation times at 20 °C and 50 °C were calculated to be 49 ± 3 and 25.6 ± 0.1 ns, respectively (fig. S4, B and C).

Distances from the residues of the hV1-S1S4 to MTSL-labeled C443 (fig. S4D), were obtained at 20 °C and 50 °C. From this data, the differences in PRE distances between 20 °C and 50 °C were calculated (ΔPRE_50 °C−20 °C_) and these values were plotted against as a function of residue number (Fig. 3E). Given the magnitudes of the temperature-dependent changes in distance, and the proclivity of PRE data to encode both structural and dynamic information, we interpret this data is a qualitative manner. The sign of the ΔPRE values indicate that a given amide proton is moving towards (negative value) or away (positive value) from MTSL-C443. The ΔPRE_50 °C−20 °C_ suggest temperature-driven changes in dynamics or conformational states in the S1-S2 loop, the S2-S3 loop, and the S4 helix C-terminus. The S1-S2 loop has complex data trends suggestive of differences in loop dynamics between temperatures. The S2-S3 loop and S4β helix C-terminus show general trends of movements away from C443. Precise interpretation on these temperature dependent PRE changes are challenging; however, we note that the S1-S2 loop, the S2-S3 loop, and the S4 C-terminus, have been implicated in proton activation (*47*), ligand binding (*20, 23*), and ligand coupling (*48*), respectively.

#### NMR-detected temperature-dependent hV1-S1S4 changes in hydration

^15^N-edited NOESY-TROSY measurements provide through-space distance information between protons up to distances of ~5 Å. NOESY data also provide a measure of solvent exposure via NOE crosspeaks between a given backbone amide proton resonance (H_N_) and the water resonance (~4.7 ppm). Since the H_α_ to H_N_ distances represent a fixed distance, taking the intensity ratios between the water-H_N_ cross-peak and the H_α_-H_N_ cross-peak (*I*_water_/*I*_Hα_) should reflect a normalized measure of solvent accessibility of a specific backbone amide resonance. Furthermore, taking the difference between the normalized ratios at high and low temperatures (ΔNOE_45 °C−20 °C_) identifies which residues are more (less) solvent exposed to the solvent at a higher (lower) temperature. With this in mind, a positive ΔNOE_45 °C−20 °C_ indicates increased solvent exposure at 45 °C. NOE-based solvent exposure between elevated and lower temperature indicate that for the vast majority (~60% of the observed residues) of hV1-S1S4, the changes in hydration are minimal. However, the S1-S2, S2-S3, and the S3-S4 loops have increased solvent exposure at higher temperatures. Similarly, the S4 helix C-terminus also become more solvent exposed at higher temperatures (Fig. 3F and fig. S5A).

#### The role of R557 in coupling between the S1-S4 domain and the pore domain

Our data show that the C-terminus of the S4 helix partially unwinds and becomes more solvent exposed. These movements include the conserved R557, as monitored from the backbone amide with PRE, RDC, chemical shift, and NOESY NMR data. A previous study that R557 in rat TRPV1 is involved in coupling the TRPV1 S1-S4 domain to channel activation in part via a cation-π interaction rearrangement between Y554 and R557 that impacts diverse activation mechanisms, including thermosensing (*48*). To probe the role of human R557-TRPV1 in temperature gating, we generated a human TRPV1-R557A mutation for electrophysiology studies. Whole-cell patch-clamp measurements of hTRPV1-R557A corroborate that this mutation inhibits TRPV1 thermosensitivity (Fig. 4, A and B) and using NMR we show that for hV1-S1S4 the R557–water NOE cross-peak is absent at 20 °C and present at 45 °C (Fig. 4C). These data indicate that R557 in the S4 helix undergoes a temperature dependent change in solvent exposure that is consistent with the ΔPRE measurements. Using NMR temperature titrations with hV1-S1S4-R557A mutant (fig. S6A), the resonance intensities retained the sigmoidal two-state model as shown in the representative plot of G548, with a Δ*H* of 19 ± 2 kcal/mol (Fig. 4D). Measuring 70 resonance intensities as before with WT, the mean Δ*H* = 19.7 ± 0.1 kcal/mol and the mean T_50_ = 39 ± 2 °C (Fig. 4E and fig. S6B). We attribute the small decrease in thermosensitivity (*ΔΔH* = 1.5 ± 0.1 kcal/mol) between the wild-type and R557A mutant of hV1-S1S4 to the loss of the cation-π interaction between R557 and Y554. If R557 couples the S1-S4 domain with the PD, then the R557A mutation should also knock-out capsaicin activation. Whole-cell patch-clamp electrophysiology of human TRPV1-R557A shows that this mutation indeed silences capsaicin activation (fig. S6C). Given the similarities of the thermosensitivity of wild-type and R557A hV1-S1S4, these data suggest that R557 functions to couple motions of the hV1-S1S4 domain and channel gating.

**Figure 4.**
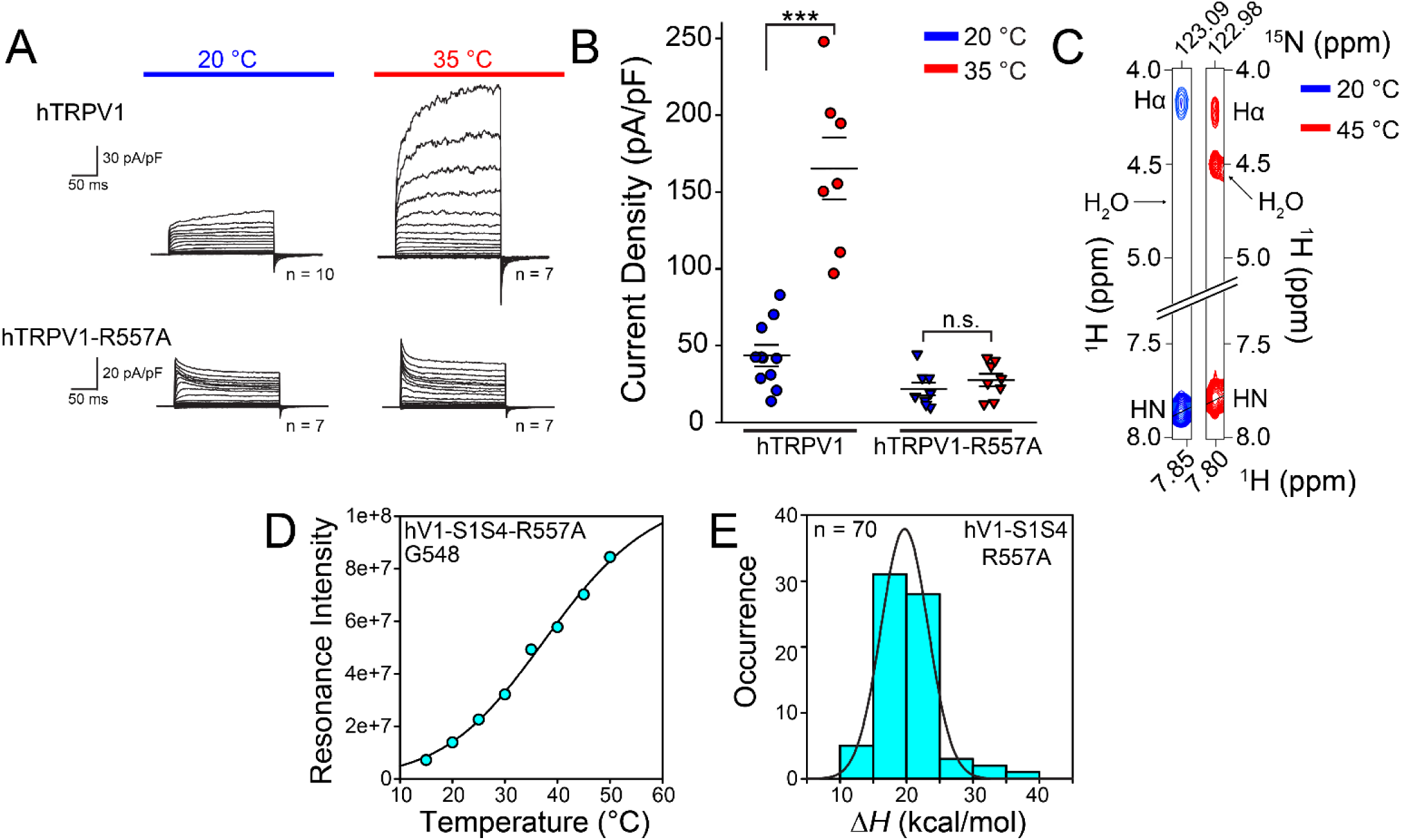
R557 is a crucial residue for coupling between the S1-S4 domain to the pore domain. (**A**) and (**B**). Whole-cell patch-clamp electrophysiology data from HEK293 cells show that the wild-type hTRPV1 is heat activated, while the hTRPV1-R557A mutant is not. (**C**) ^15^N-edited NOESY-TROSY data of R557 water resonance cross peak missing at low temperature but present at elevated temperature, suggesting that R557 amide backbone moves from a membrane embedded to a solvent accessible position in a temperature-dependent manner. (**D**) NMR temperature titration of hV1-S1S4-R557A. G548 in hV1-S1S4-R557A resonance intensities as a function of temperature show two state behavior analogous to the wild-type hV1-S1S4 domain. (**E**) A histogram and a Gaussian distribution fit of Δ*H* identify a mean ΔH = 19.7 ± 0.1 kcal/mol for the mutant hV1-S1S4-R557A (n = 70 residues). The data suggestive of a temperature-dependent conformational change in hV1-S1S4-R557A.

## Discussion

The ability to sense and respond to temperature is fundamental in biology, with a variety of temperature-sensing requirements that vary from low to high temperatures and acute (fast) to environmental (sustained) responses. Acute hot-sensing in higher organisms is sufficiently crucial that the pathway is triply redundant with three distinct heat receptors; the best studied of these is TRPV1 (*49*). Similarly, TRPV1 thermosensitivity is tuned in a highly species dependent manner that is correlated with environmental niche. For example, desert dwelling species that are exposed to sustained elevated temperatures are thought to generally have attenuated heat-sensing responses (*50*). Many factors have been shown to modulate the temperature response of TRPV1, including chemical ligands, pH, lipids, and proteins. Nonetheless, TRPV1 is intrinsically thermosensitive and the ability to integrate changes in biologically relevant temperatures as a molecular thermometer is a feature inherent to the protein (*34*). Despite its importance, the mechanism of TRPV1 thermosensing has been complicated by seemingly contradictory outcomes (*1, 2*).

In this study, we evaluate the conformational change and thermodynamics of an isolated human TRPV1 S1-S4 domain. We establish that the S1-S4 domain is stable, retains the expected membrane topology, and recapitulates anticipated capsaicin agonist binding, indicating the isolated domain resides in a biologically relevant conformation. These results are consistent with structurally homologous domains studied in isolation from other TRP (*51*), sodium (*52*), potassium (*42, 53–55*), and proton (*56*) ion channels.

Temperature dependent studies of the hV1-S1S4 monitored by CD, intrinsic tryptophan fluorescence, and solution NMR show that this domain undergoes a two-state temperature-dependent transition, with significant thermosensitivity (Δ*H* of ~20 kcal/mol). These data also indicate that the domain retains significant structure in both high and low temperature conformations. The energetics associated with this conformational change are large in comparison to other proteins. For example, the enthalpy magnitude of the hV1-S1S4 transition is about twice as large as the well-studied tetrameric hemoglobin system, despite the size of hemoglobin being about four-times larger by mass (*29*).

Putting hV1-S1S4 thermosensitivity value into the context of the full-length TRPV1 channel will require future studies. The thermosensitivity, or change in enthalpy, between temperature resting and active states of full-length TRPV1, with a consensus value is in the realm of ~100 kcal/mol (see Table S1). The value measured here for the full-length human TRPV1 heterologously expressed in mammalian cells is 98 ± 12 kcal/mol or 94 ± 8 kcal/mol, depending on the method used. One challenge with interpreting the thermosensitivity of the isolated hV1-S1S4 relative to the tetrameric TRPV1 in biological conditions is that the cooperativity between monomers during heat-sensing and activation in TRPV1 remains unclear. For hV1-S1S4, which represents ~18% of the channel by mass, could potentially contribute the majority of the thermodynamic driving force (~80%) should the Δ*H* values of the isolated hV1S1-S4 be additive in nature. On the other hand, if S1-S4 domains allosterically couple very efficiently in tetrameric channel state, then this domain might contribute as little as ~20 kcal/mol, or ~20% of the thermosensing driving force, to TRPV1 thermosensitivity.

Nonetheless, our results fit in well with the existing literature with the thermodynamic and mechanistic information about TRPV1 and thermosensing in general (*18, 32, 36, 57*). For example, studies of a TRPV1 isoform, ΔN-TRPV1, which lacks the majority of the N-terminus, including 5 of 6 ankyrin repeat domains, yet retains sensitivity to physiological changes in temperature, suggests that the core TRPV1 temperature sensing region is isolated to the transmembrane domain (*35*). Our data and a recent chimeric study of the Shaker voltage-gated potassium channel (Kv1.2) with the rat TRPV1 pore domain (S5-S6 helices, residues 575-687) suggest that both membrane domains likely contribute to thermosensing. In the chimeric study, the TRPV1 PD was sufficient to transform the chimeric Shaker channel into a thermosensing channel with a thermosensitivity of ~75 kcal/mol as estimated from the reported Q_10_ value (*36*). Further studies are required to better contextualize our thermodynamics studies with regards to the full-length channel and to decipher and quantify the cooperativity between membrane domains. If TRPV1 vanilloid ligand activation serves as a guide, concatemer studies have shown that a single capsaicin binding event can fully open the channel; on the other hand, equivalent proton activation studies, show that full activation requires all four TRPV1 proton-sensors (*58*).

Beyond thermodynamic contributions to TRPV1 thermosensitivity, there are two general views on the mechanisms of thermosensing; first, that state-dependent solvation changes, which have large thermodynamic signatures, drive thermosensing; or second, that partial unfolding as function of temperature (i.e. TRPV1 is only marginally stable) gives rise to the large thermosensitivity and drives thermosensing (*1, 29*). While mechanistic TRPV1 temperature gating details continue to emerge, our studies of hV1-S1S4 domain offer particular mechanistic insights. Our NMR studies of the hV1-S1S4 at elevated temperatures identify structural differences from the cryo-EM structures. For example, chemical shift and RDC data show that the S2, S3, and S4 helices are kinked, a feature absent in all of the current TRPV1 structures. However, S2 helical kinks are found in certain TRPV3, TRPV5, and TRPV6 structures (*59–62*). Similarly, helical distortions in S3 (*63*) and S4 are also found in other TRPV family structures. The most significant structural temperature-dependent changes in distance, hydration, and secondary structure from our studies identify three particular areas: 1) The extracellular juxtamembrane region of the S1 helix, 2) the intracellular S2-S3 loop, and 3) the intracellular S4 helix. Interestingly, these temperature-induced changes we observe are similar to those identified via molecular dynamics simulations of TRPV1 heat activation (*64*).

The extracellular S1 helix and the S1-S2 loop have been implicated in proton activation, where R455 (in rTRPV1, R456 in hTRPV1) is critical to channel function and R455K causes a gain of function (GOF) mutation (*65*). This residue in the cryo-EM structure is buried in the membrane as identified by the OPM server (*66*) and is in close proximity to lipid molecules (fig. S5C) (*21*). Moreover, this arginine is proximal to two residues that are crucial for the proton activation, V538 in S4 helix and T633 in pore helix (*47*). A recent study that engineered a RTx-sensitive TRPV3 revealed that an allosteric point mutation in the pore domain (A606V) along with mutations to the vanilloid binding pocket, is necessary for TRPV3 to be both RTx and temperature sensitive, emphasizing the role of this residue in temperature sensitivity and the allosteric modulation of the channel (*67*). In TRPV1, the equivalent residue to A606 is V596, and this residue is surrounded by R456, V538, and T633 (fig. S5C). With this region of the S1S4 domain clearly important for function, it is interesting that our temperature dependent NMR-studies identify meaningful changes in distance and hydration and suggestive that TRPV1 temperature activation might partially interface with proton activation mechanisms that are coupled to the upper pore gate.

Another area identified in our NMR studies is a difference in helicity of the S2-S3 linker relative to the cryo-EM structures. The S2-S3 linker is central to vanilloid ligand activation of TRPV1 (*23, 26, 28*) and includes Y511 and S512 that impart specificity by hydrogen bonding to the vanilloid head group of TRPV1 agonists. In the cryo-EM structures, there are helical differences between vanilloid bound and apo-TRPV1 structures. Our NMR data identify an additional helical turn in the S2-S3 linker (Fig. 1C and Fig. 3, B and C) suggestive that temperature-dependent conformational changes of this domain interface with features key to vanilloid activation.

A common feature of TRPV family structures is a stretch of 3_10_ helix at the intracellular side of the S4 helix. Interestingly, this is not present in our NMR investigations of the hV1-S1S4, instead we see a kinked α-helix where the transition between α-and 3_10_-helix is found in most TRPV structures. In addition, we see in the RDC data that there is a loss of one helical turn at the S4 C-terminus (Fig. 3C). This loss of helix is coupled with a general movement of the S4 helix towards C443 in the S1 helix, as identified in PRE measurements (Fig. 3E), indicating a concerted movement towards the center of the membrane plane. These changes in secondary structure and distance are concomitant with an increase in hydration of the S4 C-terminus (Fig. 3F). Similar to the loss of helicity at the C-terminus end of the S4 helix in the hV1-S1S4, the unfurling of the bottom of the S4 helix in the activated state is found in TRPV1, TRPV2 and TRPM8 (fig. S7, A to C). This helix unfurling is also shown in the apparent TRPM8 ligand-gating mechanism (*68*). Upon an agonist binding, the S4 helix undergoes changes in secondary structure from α-to 3_10_ helix, starting around V839, pushing the membrane-bound R850 out the membrane (fig. S7C). This R850 in TRPM8 is equivalent to R557 in TRPV1, as the alignment of the S1-S4 domains of TRPV1 and TRPM8 shows that R557 in TRPV1 and R850 in TRPM8 are located in the same region in the membrane (fig. S7D).

The C-terminal S4 region is functionally important in coupling the S1-S4 domain to the pore domain (*48*). Our data identify that R557 in S4b, which is membrane bound in the cryo-EM structures and at 20 °C in our NOE-detected solvation studies, becomes water accessible at elevated temperatures. This residue is well-conserved, and mutation to lysine (R557K) results in a GOF mutation (*48*). The TRPV1 cryo-EM structures have provided molecular insight into the role of R557 in ligand-activated channel coupling. In the apo structure R557 forms a cation-π interaction with Y554 and a lipid separates R557 from interacting with E570 in the S4-S5 linker (Fig. 5A). Vanilloid binding releases the bound lipid separating R557 from E570 allowing for a switch in R557 conformation that leads to channel activation (Fig. 5A) (*21*). Our data indicate that heat causes similar conformational changes that would result in the disruption of the cation-π interaction between Y554 and R557, as R557 changes helicity and becomes more solvent accessible at elevated temperatures (Fig. 5B). Presumably, these changes could couple analogously to vanilloid binding to gate the pore domain. To probe this, we generated an R557A mutant, and electrophysiology studies show that it has both impaired heat and capsaicin sensitivities (fig. S6C), consistent with the disruption of S1-S4 domain coupling to gating. The R557A mutation was also incorporated into the isolated hV1-S1S4 and the NMR data show that it retains structure, and only modestly impacts thermosensitivity by ~1.5 kcal/mol relative to the wild-type (fig. S6B). Presumably this slight loss of thermosensitivity is from the oblation of the Y554-R557 cation-π interaction. In TRPV4, the residues equivalent to Y554 and R557 in TRPV1 (Y591 and R594) have also been implicated in ligand activation, and mutations in the TRPV4 S3 and S4 helices also impacts heat activation (*69*). Taken together these data suggest that R557 functions to couple both heat (Fig. 4, A and B) and ligand (fig. S6C) stimuli to the pore domain in gating and is in agreement with a mechanism proposed by Cheng, Julius, and coworkers (*21*). We note that R557 is not included in the construct used in a recent study that swapped the TRPV1 PD to the temperature-insensitive Shaker channel (fig. S6D) (*36*), suggesting that in the context of the full-length TRPV1 channel, the PD is not exclusively sufficient to produce thermal activation. Our studies suggest that interplay between these membrane domains are required for heat activation.

**Figure 5.**
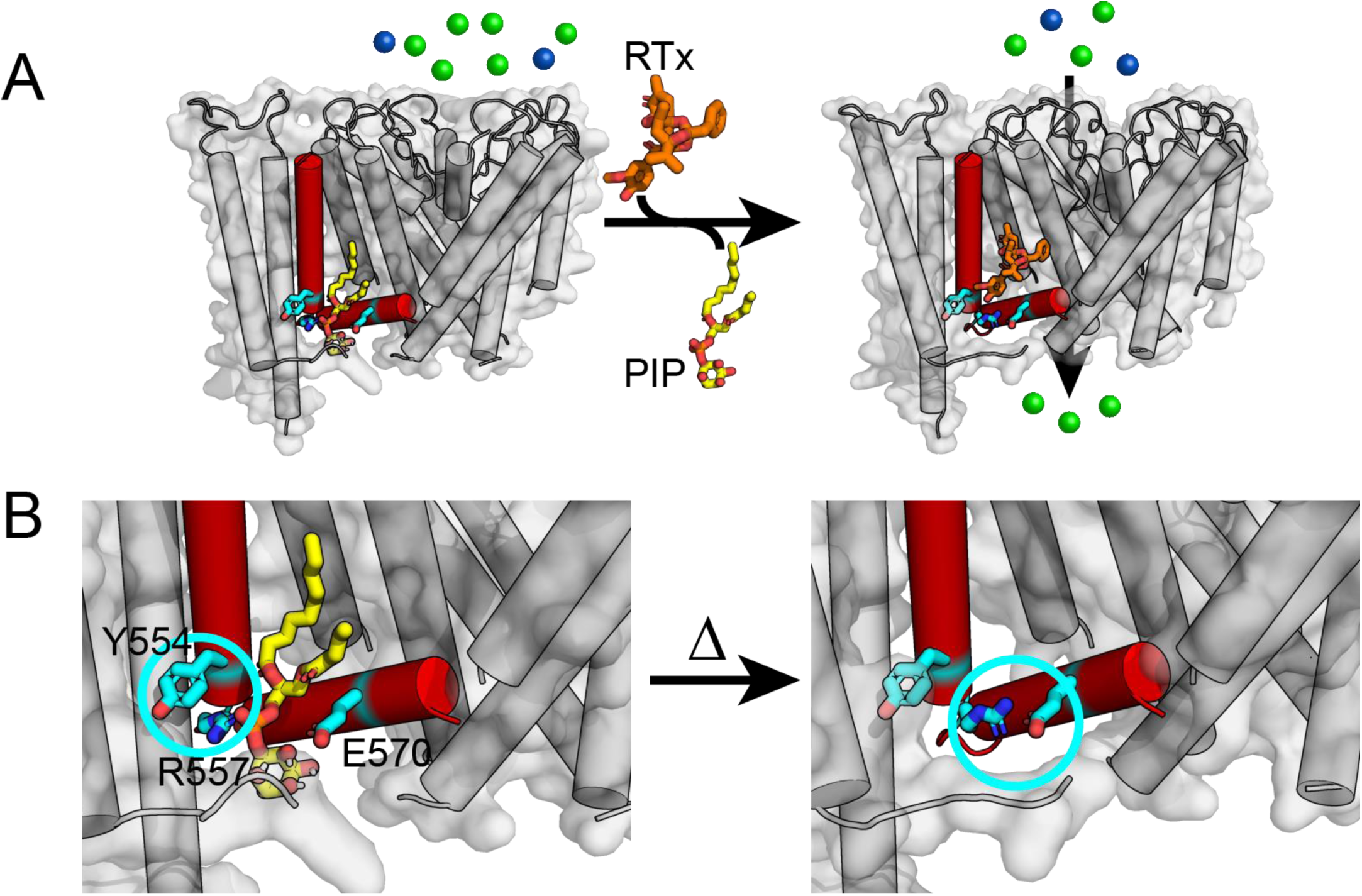
A proposed heat-sensing mechanism of TRPV1 involves the interaction between the S4 helix and the S4-S5 linker. (**A**) Structure-based ligand-gated mechanism of TRPV1 is shown in cartoon (PDB ID: 5IRZ and 5IRX). In the resting state, an endogenous lipid (shown in yellow) occludes the canonical vanilloid binding pocket, perturbing the interaction between the R557 in S4 and E570 in S4-S5 linker. As a vanilloid ligand, resiniferatoxin (RTx, orange), binds the S1-S4 domain, the cation-π interaction between R557 and Y554 is disrupted and R557 changes conformation to interact with E570, opening the lower gate. For visual clarity, S3 helix is omitted, and only one S1-S4 domain is shown. (**B**) The close-up views that show the interaction between Y554 and R557 in S4 helix in the resting state (left, highlighted in cyan circle), and the interaction between R557 and E570 in S4-S5 linker (right, shown in cyan circle). Our data suggest that the C-terminal end of the S4 helix undergoes heat-induced movements in this region and lose the helicity. This model is analogous to the ligand activation mechanism and consistent with a heat-activation mechanism proposed by Cheng, Julius, and coworkers (*21*).

Overall, our study provides new insights into the role of the S1-S4 domain in TRPV1 gating. We present the first direct binding measurements of capsaicin to the S1-S4 domain and validate the identity of key residues exhibit specific capsaicin binding. Utilizing various techniques, we show that the S1-S4 domain undergoes a temperature-dependent conformational change with a relatively large magnitude Δ*H* (thermosensitivity). Mechanistic studies suggest that there is overlap between regions of the S1-S4 domain that have been previously implicated in proton and ligand activation and the conformational changes identified in our study. Taken together our data indicate that the S1-S4 domain contributes to TRPV1 thermosensitivity.

## Materials and Methods

### Human TRPV1-sensing domain (hV1-S1S4) protein expression and purification

The hV1-S1S4 (S1-S4, including Pre-S1) was expressed from a synthetic gene engineered from ATUM that encodes the hV1-S1S4, a 141-residue structural domain with an identical amino acid sequence to human TRPV1. The optimized gene was modified to include an N-terminal 10×His tag fused to a thrombin cleavage site subcloned into a pET16b vector. The expressed protein includes residues Pro418-Gly558 from the human TRPV1 channel.

We optimized the fusion hV1-S1S4 expression using *Escherichia coli* strain BL21 (DE3) competent cells. Starter cultures were initiated with 1 transformed colony inoculated in a culture tube containing 3 mL LB and 0.1 mg/mL ampicillin (3 μL of 100 mg/mL stock) (Sigma Aldrich) and incubated for ca. 7 h at 37 °C. 6 mL of starter culture was used to inoculate 1 L of M9 minimal media (12.8 g Na_2_HPO_4_·7H_2_O (Sigma Aldrich), 3 g KH_2_PO_4_ (Thermo Fisher Scientific), 0.5 g NaCl (Thermo Fisher Scientific), 1 mM MgSO_4_·7H_2_O (Sigma Aldrich), 0.1 mM CaCl_2_·2H_2_O (Sigma Aldrich), 4% D-glucose (w/v) (Sigma Aldrich), 1× MEM Vitamin solution (100× solution, Corning)) with 1 g ^15^NH_4_Cl (Cambridge Isotope Laboratories) as the sole nitrogen source. The cells were grown at 18 °C and the hV1-S1S4 expression was induced for 48 h by addition of 0.1 mM isopropyl β-D-1-thiogalactopyranoside (IPTG, Research Products International Corp.) at an OD_600nm_ of 0.5-0.6. After 48 h, the cells were harvested by centrifugation at 6,000 ×*g* for 20 min at 4 °C resulting in a cell pellet average of ca. 2.5 g per liter of M9.

The cell pellet was resuspended in 25 mL lysis buffer (75 mM Tris-HCl (Thermo Fisher Scientific), 300 mM NaCl (Thermo Fisher Scientific), 0.5 mM EDTA (Sigma Aldrich), pH 7.7) with 0.01 mg of lysozyme (Sigma Aldrich), 0.01 mg DNase (Sigma Aldrich), 0.01 mg RNase (Sigma Aldrich), 1.25 mg PMSF (Sigma Aldrich), 0.05 mM magnesium acetate (Sigma Aldrich) and 2 mM β-mercaptoethanol (β-ME, Sigma Aldrich). This lysis solution was tumbled at room temperature for 30 min. The mixture was sonicated using an S-4000 Ultrasonic Processor (Qsonica) at 4 °C for 7.5 min at 55% power and 50% duty cycle of 5 sec on and 3 sec off. The resulting whole cell lysate was then tumbled with N,N-dimethyl-N-dodecylglycine (Empigen BB Detergent, BOC Sciences) (3% v/v) for 1 h at 4 °C to solubilize and extract the hV1-S1S4 into micelles. After an hour, the lysate was centrifuged at 38,500 ×*g* for 30 min at 4 °C to pellet non-solubilized cellular debris. The supernatant was collected and tumbled with 1 mL of preequilibrated Ni(II)-NTA Superflow (Qiagen) resin for 1-1.5 h at 4 °C.

The hV1-S1S4-bound resin was packed in a gravity column to a column volume ~ 1 mL and washed with 20 column volumes of Buffer A (40 mM HEPES (Sigma Aldrich), 300 mM NaCl (Thermo Fisher Scientific), 2 mM β-ME (Sigma Aldrich) pH 7.8) with 1.5% (v/v) Empigen detergent (BOC Sciences). The resin was washed with 20 column volumes of wash buffer (40 mM HEPES, 300 mM NaCl, 60 mM Imidazole (Sigma Aldrich), 1.5% (v/v) Empigen, 2 mM β-ME, pH 7.8). With the hV1-S1S4 bound to the resin, the detergent was exchanged into the desirable membrane mimic for further studies with 15 column volumes of rinse buffer (25 mM Na_2_HPO_4_ (Thermo Fisher Scientific), 0.05% (w/v) lyso-palmitoylphosphatidylglycerol (LPPG, Anatrace), 2 mM β-ME, pH 7.8). Finally, the hV1-S1S4 was eluted with 15 column volumes of elution buffer (25 mM Na_2_HPO_4_, 300 mM Imidazole, 0.1% (w/v) LPPG, 2 mM BME, pH 7.8).

A variety of membrane mimics were screened by ^15^N-TROSY-HSQC NMR, including the following: DPC (0.5% (w/v)), TDPC (0.5% (w/v)), LMPC (0.2% (w/v)), LMPG (0.2% (w/v)), LPPC(0.2% (w/v)), and LPPG (0.2% (w/v)) (Fig. S1). Spectra of all tested samples were collected at both 20 °C and 50 °C, and the spectral qualities were compared. The hV1-S1S4 purified in LPPG micelles produced the most well-resolved and dispersed spectra with the appearance of the expected number of resonances and was therefore the membrane mimic chosen for the resulting studies.

After Ni-NTA purification the hV1-S1S4 was buffer exchanged to thrombin cleavage buffer (25 mM Na_2_HPO_4_, 150 mM NaCl, pH 7.8) using an Amicon Ultra centrifugal ultrafiltration unit (Milipore, 10 kDa cutoff). The concentration of protein was determined using a BCA kit (Thermo Fisher) and 3U thrombin (Novagen) was added for every 1 mg protein. The thrombin-protein cleavage reaction was incubated at room temperature (ca. 23 °C) for 24 h. After the reaction was completed, this mixture was passed over Ni-NTA resin with the flow through containing the thrombin-cleaved hV1-S1S4.

Gel filtration of the cleaved protein was carried out by concentrating to ca. 0.5 mL in NMR buffer (25 mM Na_2_HPO_4_, 0.5 mM EDTA, pH 6.5), and was eluted with a 1 CV of a Superdex 200 (GE Healthcare Life Sciences) column. The fractions with high A_280_ readings were analyzed with SDS-PAGE. The purified sample was concentrated to a desired volume. After purification, the average yield of pure hV1-S1S4 per liter of M9 medium was approximately 1.5 mg.

### Validation of the hV1-S1S4 Identity

To confirm the identity of purified hV1-S1S4, the SDS-PAGE band was cut and sent to the MS Bioworks, which performed liquid chromatography tandem mass spectrometry (LC-MS/MS). The result revealed that our purified hV1-S1S4 matches the hV1-S1S4 amino acid sequence we provided with 51% coverage (fig. S1B).

### MTSL Site Directed Spin Labeling

For paramagnetic relaxation enhancements (PRE) maleimide chemistry was used to label the hV1-S1S4 with a nitroxide spin label MTSL (*S*-(1-oxyl-2,2,5,5-tetramethyl-2,5-dihydro-1H-pyrrol-3-yl)methyl methanesulfonothioate, Santa Cruz Biotech). Only the WT construct containing a lone cysteine in S1 α-helix (C443) was used for PRE. For these reactions, the hV1-S1S4 was prepared as above, except before thrombin cleavage the volume of purified hV1-S1S4 was adjusted to 0.5 mL in 25 mM sodium phosphate buffer at pH 7.2. To maximize nitroxide labeling efficiency, purified hV1-S1S4 was incubated and tumbled with 2.5 mM dithiothreitol (DTT, A Geno Technology, Inc.) for 1-2 h at room temperature to reduce the cysteine sulfhydryl. After the reaction with DTT, a 10:1 MTSL (stock solubilized in DMSO) to protein mole ratio amount of nitroxide spin label was solubilized in DMSO and incubated for 3 h at 37 °C. The reaction was continued at room temperature overnight and was buffer exchanged to 25 mM Na_2_HPO_4_, pH 7.8. The resulting MTSL-hV1-S1S4 was rebound to Ni-NTA resin and incubated for 1 h at room temperature. MTSL was removed by washing the resin with at least 20 column volumes of rinse buffer (25 mM Na_2_HPO_4_, 0.05% (w/v) LPPG, pH 7.8). MTSL labeled-hV1-S1S4 was eluted with 5 column volumes of elution buffer (25 mM Na_2_HPO_4_, 300 mM Imidazole, 0.1% (w/v) LPPG, pH 7.8). After this step, the purification follows as described above with thrombin cleavage and SEC.

### Nuclear Magnetic Resonance Spectroscopy

All NMR experiments were recorded with a Bruker 850 MHz ^1^H spectrometer and Avance III HD console equipped with a 5 mm TCI CryoProbe. All samples included 4 μL D_2_O with a total volume of 180 μL in a 3 mm NMR tube. The temperature for these experiments was calibrated using 99% ^2^H methanol where the difference in chemical shifts (ppm) arising from the two resonances is used as a temperature standard with the ranges of 178 K and 330 K according to *T*[*K*] = 409.0 − 36.54Δ*δ* − 21.85(Δ*δ*)^2^ where Δ*δ* is the difference in Hz between the two peaks (*70*).

#### NMR Backbone Resonance Assignment

Experiments used for backbone resonance assignment of the hV1-S1S4 were carried out at 45 °C with a 900 µM uniformly ^15^N, ^13^C labeled sample in a Bruker shaped NMR tube (Part Number Z106898) with 4% D_2_O (v/v). TROSY versions of HSQC, HNCA, HNCOCA, HNCO, HNCACO, HNCACB, and CBCACONH as well as non-TROSY 4D HNCACO and 4D HNCOCA experiments were utilized, and an ^15^N-edited HSQC-NOESY with a 90 ms mixing time (τ) was used for sequential assignments; table S1 details specific experimental parameters. The nonuniformly sampled data were reconstructed with qMDD and processed in nmrPipe, with analysis and resonance assignment carried out using CcpNMR (*71*) software (fig. S1, E to J). The resonance assignments at 45 °C are deposited in the Biological Magnetic Resonance Bank (BMRB entry 27029).

#### Secondary Structure Calculations and Comparisons with rTRPV1 cryo-EM structures

The secondary structure of the hV1-S1S4 at 45 °C was determined using TALOS-N (*39*) where the secondary structure was plotted as probability of secondary structure vs residue number. The secondary structure was then plotted in Aline (*72*) alongside rTRPV1 structures determined by cryo-EM. The secondary structure for each of the rTRPV1 structures were specifically determined by analyzing the PDB codes deposited in the Protein Data Bank for requisition numbers 5IRZ, 5IRX, and 5IS0.

#### ^1^H-^15^N TROSY-HSQC-detected Capsaicin Titrations

For the titration with capsaicin (Corden Pharma), stock capsaicin was prepared in 195 proof ethanol (Thermo Scientific), and the mole percent of capsaicin was calculated by the equation below:

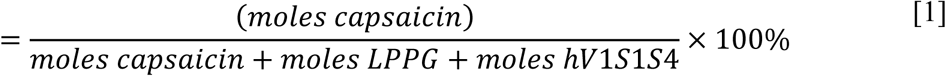

Using 66 μM hV1-S1S4 in 1.67% LPPG (w/v), ^1^H-^15^N TROSY-HSQC spectra of for capsaicin titration were collected at 37 °C, with 32 transients of 2048 direct points and 128 indirect points. The hV1-S1S4 resonance assignments at 45 °C were mapped to 37 °C and used to identify residues that specifically interact with capsaicin (Fig. 1E). The mole% capsaicin used were 0.001%, 0.002%, 0.005%, 0.012%, 0.015%, 0.025%, 0.075%, 0.125%, 0.225% and 0.723% (Fig. 1E). These spectra were processed with nmrPipe (*73*) and the resonances were analyzed in CcpNmr (*71*). Chemical shift perturbations induced by capsaicin were calculated by the equation below:

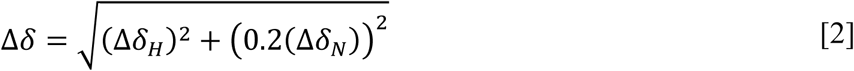

where Δδ_H_ is the difference in chemical shift in the proton dimension and Δδ_N_ is the difference in chemical shift in the nitrogen dimension, and the Δδ_N_ was multiplied by a scaling factor of 0.2 typically used for nitrogen nuclei (*74*). The capsaicin-dependent chemical shift perturbations were plotted as a function of the mole% of added capsaicin and fit to a single site binding model derived from the Langmuir adsorption model commonly used to describe binding in proteins:

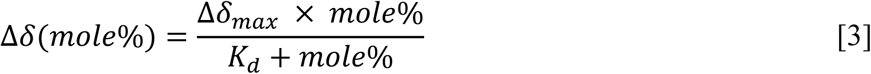

where *K*_d_ is the dissociation constant and Δδ_max_ is the maximal value of change in chemical shift from perturbation. Using the amide backbone assignment of the hV1-S1S4, the chemical shift perturbations of 72 residues were analyzed and the *K*_d_ values for each residue were calculated using *MATLAB R2016a*fit to equation [3].

#### ^1^H-^15^N TROSY-HSQC-detected Temperature Titrations

The ^1^H-^15^N TROSY-HSQC experiments of the purified hV1-S1S4 were recorded from 15 °C to 50 °C, at 5 °C increments with all other parameters fixed. Specific types of buffers exhibit pH changes at different temperatures, an effect that could convolute our analyses. To ensure that the chemical shift perturbations were caused exclusively by temperature changes and not from pH changes of the buffer, ^1^H-^15^N-TROSY-HSQC spectra at two different pH values, 6.5 and 6.4, were collected at 37 °C (fig. S3G). These two pH values were chosen based on empirical pH changes measured with our NMR buffer as a function of temperature (Table S2). No chemical shift perturbations were detected allowing direct analysis for their dependence on temperature dependent resonance changes.

The resonance intensities of several hV1-S1S4 residues were plotted against the temperature, and fit to a sigmoidal two-state model using *MATLAB* scripts relying on the *nlinfit* function with the following equation derived from a Van’t Hoff model:

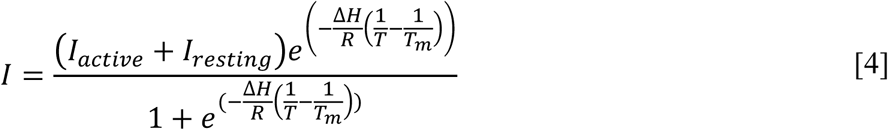

From this equation, the changes in enthalpies (Δ*H*) of 71 residues for WT and 70 residues for the hV1-S1S4-R557A mutant residues were obtained. For the WT, three distinct samples were used for the experiments, yielding three sets of the Δ*H* values per residue. From the enthalpies, a histogram with bins of 5 kcal/mol was fit to a Gaussian function,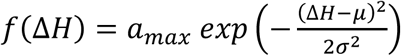, in *MATLAB* for both WTand R557A, where μ and σ are the ensemble average enthalpy and variance respectively.

#### <u>T</u>ROSY for <u>r</u>ot<u>a</u>tional <u>c</u>orrelation <u>t</u>imes (TRACT) measurements

To accurately quantify the PRE-derived distances between individual indole amine (W549 and W427) residues and the MTSL-labeled C443, accurate rotational correlation times (τ_c_) are needed at 20 °C and 50 °C. To approximate τ_c_, TRACT (*75*) was collected over a range of time delays from τ = 0 to 100 ms with 4 ms intervals. The pulse program that was used for these experiments was adapted from Lee, D et al (*75*). The data were processed and integrated in nmrPipe over a range of 9.6 ppm to 10 ppm, to which a linear baseline correction was applied with the nmrPipe function *BASE* and a defined node list in the noise (fig. S4B). The range that was integrated was selected as it encompassed the W549 indole amine resonance from the S4 transmembrane helix, and only this was integrated since integration over the entire amino region can artificially inflate the effective correlation time (*76*). The integrated data was fit using *MATLAB R2015a* with a monoexponential decay function, *f*(*τ*) = *Ae*^−*βτ*^, where *A* is a normalized maximum, τ is the time delay, and β is the relaxation decay constant in Hz. The relaxation decays for the α and β state of exclusively the W549 indole amine resonance were used to calculate the approximate effective rotational correlation time (τ_c_) due to the rigid body assumption as described previously by Lee, D et al (*75*).

#### Calculation of Amide Proton Temperature Coefficients

From the NMR temperature titration from 15 °C to 50 °C (288 K to 323 K), the amide proton chemical shifts (δ_HN_) of assigned resonances at corresponding temperatures were plotted in the same temperature range. Each plot was fit to the linear function, *y* = *mx* + *b*, where the slope is the temperature coefficient, Δδ_HN_/ΔT (ppb/K).

#### Paramagnetic relaxation enhancement (PRE) measurements

Paramagnetic (MTSL labeled) and diamagnetic (without the MTSL label) proton transverse relaxation (R_2_) measurements were recorded using a TROSY-HSQC pulse program modified to include a relaxation delay before acquisition as described by Clore and coworkers (*77*). Four relaxation delays (0, 4, 12, 24 ms) were recorded for the matched paramagnetic and diamagnetic samples at 20 °C (196 scans) and 50 °C (80 scans) with 2048 direct point and 128 indirect points. The spectra were processed using nmrPipe with a Gaussian apodization function, which does not affect the calculated distance to the free electron spin (*78*). The resonance intensity values were obtained using CcpNmr and transverse relaxation rates of the paramagnetic state, R_2_^eff^, and the diamagnetic state, R_2_, were extracted by fitting the data with a monoexponential decay function, *y* = *ae*^−*bx*^ (fig. S4E). The paramagnetic rate enhancements, Γ_2_, were calculated by subtracting R_2_ from R_2_^eff^, using the relationship of the electron spin-enhanced transverse relaxation rate due to the spin label (R_2_^eff^) is a sum of the intrinsic transverse relaxation rate (R_2_) and the contribution from the electron (Γ_2_):

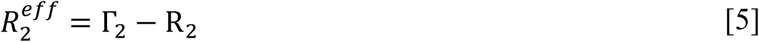

The errors for R_2_^eff^ and R_2_ were extracted from fitting errors, and the error of δΓ_2_ was calculated using standard the propagation of error:

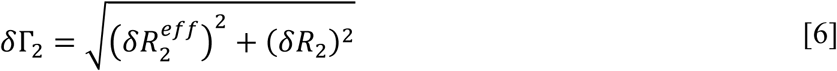

The distance restraints from MTSL-labeled C443 to residues in proximity were calculated by the following relationship:

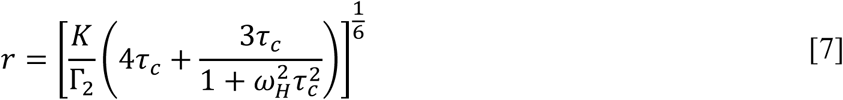

where *τ*_*c*_ is the effective rotational correlation time, ω_H_ is the Larmor frequency for the nuclear spin (protons), and K is 1.23 × 10^−32^ cm^6^ s^−2^ a constant related to the gyromagnetic ratio (γ), spin quantum number (S), electronic g factor (*g*), and the Bohr magneton (β) according to 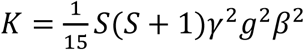.

Three separate paramagnetic (MTSL labeled) and diamagnetic (non-MTSL labeled) samples were prepared. Transverse relaxation rates of states were obtained, and using various combinations, 9 paramagnetic enhancement rates were calculated, generating 9 distance restraints per residue. Distances calculated to be higher than 27 Å were disregarded as this is beyond the reliable distance calculation for the atoms involved due to its r^−6^ dependence. The distances at 20 °C and 50 °C were averaged, and the temperature dependent change in distances (ΔPRE) were calculated by subtracting 20 °C (resting state, initial) from 50 °C (active state, final) (ΔPRE_50 °C – 20 °C_). We interpreted that the residues with negative ΔPREs are moving closer to MTSL-C443, and the residues with positive ΔPREs are moving away from MTSL-C443. The errors of distance restraints were the standard error of the mean (SEM).

#### NOESY at two different temperatures

^15^N-edited TROSY-HSQC-NOESY spectra of the hV1-S1S4 were collected with 96 scans and 48 scans with the mixing time of 90 ms at 20 °C and 45 °C, respectively. Using the amide resonance assignments and temperature titration resonances, amide NOESY crosspeaks were assigned to corresponding Hα and water cross-peaks. Using peak volumes, the ratio (*I*) between the water and Hα cross-peaks were calculated by dividing the water cross-peak by Hα cross-peak at both temperatures. Then the differences of *I*_water_/*I*_Hα_ at 45 °C and 20 °C (ΔNOE_45 °C – 20 °C_) were calculated for all well resolved resonances to observe if the hV1-S1S4 undergoes changes in hydration as a function of temperature. The ΔNOE_45 °C – 20 °C_ was plotted against the residue number and the cutoff value of 0.5 was used to evaluate the changes in hydration per residue.

#### Deuterium/hydrogen exchange factor analysis of the hV1-S1S4 at 50 °C

The H_2_O/D_2_O exchange measurement for the hV1-S1S4 was carried out at 50 °C. To vary the concentration of D_2_O in the sample, the NMR buffer was prepared in D_2_O. This was done by lyophilizing 500 μL of NMR buffer and resuspending it in 500 μL of D_2_O. To make 10% D_2_O/90% H_2_O, 18 uL of NMR buffer in D_2_O was added to the sample, and the sample was incubated for approximately 12 hours to ensure that the exchange between the proton and deuterium is completed. ^1^H-^15^N TROSY-HSQC was collected as described above with 32 scans. For the higher concentration of D_2_O titration points, the sample with 10% D_2_O/90% H_2_O was concentrated down to approximately 100 μL, and 80 μL of the NMR buffer in D_2_O was added to the concentrated sample to make a 40% D_2_O/60% H_2_O sample. For the further D_2_O points, the same method was used.

The resulting data was analyzed with a method developed by Opella et al. (*79*). The relative peak volumes of resonances were normalized to those from the 2.2% D_2_O spectrum. The mole fraction of water (χ_water_) was calculated for each data point and plotted against the normalized peak volumes. The decay in resonance intensity as a function of χ_water_ is linearly dependent on the exchange factor *m*. All D/H assigned exchange factors were plotted against the residue number for further analysis.

#### Residual Dipolar Coupling (RDC)

A neutral 3.8% polyacrylamide gel comprising 3.8% (w/v) copolymer was generated using acrylamide (stock 40% w/v, Sigma-Aldrich), bis-acrylamide (stock 2%, Sigma-Aldrich), 10% APS, and 4 μL of TEMED in a casting solution of 10 mL of 250 mM imidazole at pH 6.5. These gels were cast in a 3D printed plastic block with a custom 6 mm inner diameter polytetrafluoroethylene (PTFE, Teflon) tube insertion. Prior to adding TEMED, the solution was filtered using 0.22 µm filter (Millipore) to eliminate any polymerized acrylamide. The polymerization reaction was carried out for 24 hours. Once polymerization was complete, the gel was soaked in ddH_2_O for ~12 h initially, then it was dialyzed in the NMR buffer for 48 hours with a gentle nutating at room temperature; with buffer changes every 12-18 h. Once the dialysis was complete, the gel was cut to 1.7 mm length; the length was empirically chosen to span the probe coil within a 5 mm NMR tube. The cut gel was partially dehydrated by incubating at 37 °C for 3.5 h, which was then soaked in 1 mL of a purified 177 μM hV1-S1S4 sample for approximately 48 h. The concentration of the protein solution outside of the gel was measured every 24 h to monitor absorption into the gel. Once concentration plateaued, the protein-soaked gel was stretched into a 5 mm NMR tube with an inner diameter of 4.4 mm using Tygon tubing attached to the NMR tube and a syringe, the difference in gel and tube diameter stretched the gel to ~4 mm. Residual dipolar couplings were measured using <u>A</u>mide <u>R</u>DC’s by <u>T</u>ROSY <u>S</u>pectroscop<u>y</u> (ARTSY) by taking the ratio of assigned amino acid intensities from attenuated spectra and reference spectra (*80*). The dipolar couplings were then plotted as a function of residue number and fit according to the following equation:

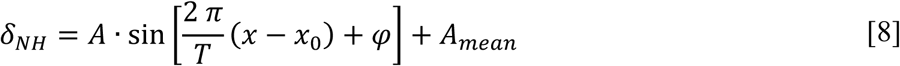

where *T* is the period of the wave, *x* is the amino acid identity, *x0* is the first amino acid to be fit in a given range, *A*_*mean*_ is the average of the couplings in a given range, and *A* represents a function of the alignment tensor according to:

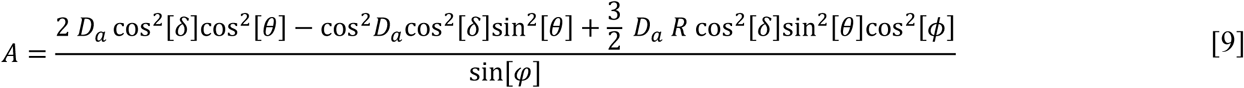

where *R* and *D*_*a*_ are the rhombicity and axial symmetry of the alignment tensor, respectively. The angles θ and ϕ correspond to the amide bond vector angles with respect to the Y- and Z-axes, and δ the angle that the amide bond vector makes with the chain axis.(*81*) The first equation is a good approximation for ideal α-helices and 3_10_ -helices as previously described (*81, 82*) RMSE was calculated according to:

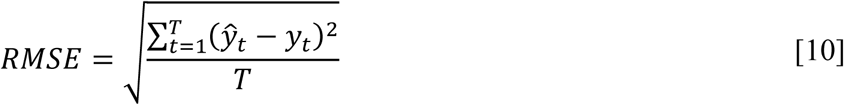

where *ŷ*_*t*_ corresponds to predicted values of *y*_*t*_ over *T* observations.

### Far-ultraviolet circular dichroism (Far-UV CD)

Far-UV CD measurements were carried out on a 0.2 mg/ml hV1-S1S4 sample in 0.1% (w/v) LPPG micelles in 200 µL in 25 mM Na_2_PO_4_ buffer at pH 6.5. CD spectra were obtained with a Jasco J-710 spectropolarimeter using a 1.0 mm path length cell. The temperature was controlled with a Peltier device (Jasco PTC-424S). All experiments were measured from 250 nm to 190 nm and 5 scans from 10 °C to 57 C in 1 °C increments. The blank (25 mM Na_2_HPO_4_, pH 6.5, containing 0.1% (w/v) LPPG) measurement values were subtracted from the protein measurement values, and the units of CD values were converted into mean residue ellipticity using the equation:

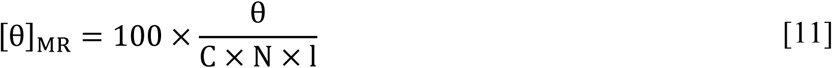

where [θ]_MR_ is the mean residue ellipticity, θ is ellipticity in millidegrees, C is the concentration of protein in molar (M), N is the number of amino acid residues of protein, and l is the path length in centimeters. The hV1-S1S4 CD values at 222 nm were plotted against temperature (Fig. 2B).

### Temperature study using intrinsic tryptophan fluorescence measurement

Fluorescence emission spectra were measured on a QM-4/2005SE Spectrofluorometer (PTI, NJ) using 295 nm excitation to minimize contributions from tyrosine residues (*38*). The temperature was controlled over a range of ca. 7 °C – 50 °C using a water circulation system and a calibrated thermocouple was used to measure the temperature of the sample inside the cuvette. To account for light scattering contributions, a sample of LPPG micelles and buffer was used as a blank and subtracted from the measurements containing hV1-S1S4. The blank signals are relatively small and their magnitude and position do not change with temperature (fig. S4G). In contrast, for the hV1-S1S4 samples, increasing temperature resulted in both a decrease in fluorescence intensity and a spectral shift to higher wavelengths. We characterize fluorescence spectral shifts in terms of the average emission wavelength, ⟨λ⟩:

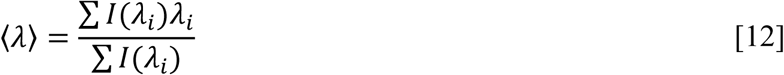

where *I*(*λ*)_*i*_ is the intensity measured at wavelength *λ*_*i*_. This quantity is more precise and less sensitive to instrumental noise than the peak maximum (λ_max_) because the calculation reflects data from the entire spectrum rather than a single point (*83*).

### Functional Measurements

#### Cell culture

HEK-293 cells (ATCC cell line CRL-1573) were cultured in 35 mm dishes at 37 °C in DMEM with 10% fetal bovine serum, 2 mM L-glutamine, 4.5 mg ml^−1^ glucose, and 100 mg ml^−1^ each of penicillin and streptomycin in the presence of 5% CO_2_. All reagents were obtained from Life Technologies.

#### Plasmid and mammalian cell transfection

The full-length human TRPV1 gene was subcloned into a pIRES-2 plasmid also containing the EGFP gene. This construct produces bicistronic mRNA containing an internal ribosome entry site (IRES) between the two genes, allowing for the independent translation of TRPV1 and the EGFP reporter. TRPV1-Arg557Ala was generated in the same plasmid using standard site-directed mutagenesis. Cells were transiently transfected with 0.4 µg DNA using FuGENE 6 transfection reagent (Promega) in a 1:3 µg DNA:µL FuGENE ratio 48 h before electrophysiology measurements were performed.

#### Electrophysiology

Transfected cells were released from the culture dish surface by briefly exposing them to 0.25% trypsin-EDTA (Thermo) and resuspending in supplemented DMEM. Cells were plated on glass coverslips and incubated for 1-2 h at 37 °C. Whole-cell voltage-clamp measurements were performed with an Axopatch 200B amplifier and pClamp 10.3 software (Molecular Devices). Data was acquired at 2 kHz, filtered at 1 kHz, and digitized using a Digidata 1440a digitizer (Molecular Devices). Patch pipettes were pulled from borosilicate glass capillaries (World Precision Instruments) using a P-2000 laser puller (Sutter Instruments) and heat polished with a MF-830 microforge (Narashige), resulting in resistances of 2-5 MΩ. A reference electrode was inserted into a salt bridge composed of 2% agar in extracellular solution. Glass coverslips plated with cells were placed in a chamber and covered with extracellular solution containing 132 mM NaCl, 5 mM KCl, 1 mM MgCl_2_, 2 mM CaCl_2_, 10 mM HEPES, and 5 mM glucose. The pH of the solution was adjusted to 7.4 using NaOH and osmolality was adjusted to 310 mOsm using sucrose. Pipette solution contained 315 mM K^+^ gluconate, 5 mM KCl, 1 mM MgCl_2_, 5 mM EGTA, and 10 mM HEPES, with pH adjusted to 7.2 using KOH and osmolality adjusted to 300 mOsm using sucrose. Osmolality was measured using a Vapro 5600 vapor pressure osmometer (Wescor). Temperature was controlled by perfusing preheated or cooled extracellular solution using an HCPC perfusion system and HCT-10 temperature controller (ALA Scientific), which heats or cools solution by supplying a specified voltage to a Peltier device through which perfusion solution flows. Temperature was calibrated by measuring the temperature of the solution exiting the HCPC at a given voltage. Currents were recorded at +60 mV and normalized to cell membrane capacitance.

## Supplementary Materials

### Materials and Methods

Details of the thermodynamic framework for interpreting temperature-dependent studies.

**Fig. S1.** The expression and NMR amide backbone assignment of the hV1-S1S4.

**Fig. S2.** Temperature-dependent data of TRPV1 and hV1-S1S4.

**Fig. S3.** The thermodynamic analyses of the hV1-S1S4 using various biophysical methods.

**Fig. S4.** Additional data supporting the thermodynamic analyses of the hV1-S1S4 using various biophysical methods.

**Fig. S5.** Supporting data for RDC and PRE measurements.

**Fig. S6.** The hTRPV1-R557A mutant becomes insensitive to both temperature and capsaicin, but the hV1-S1S4-R557A retains the temperature sensitivity.

**Fig. S7.** Structural examples of S4 helix motions in TRP channel activation.

**Supplementary Table 1.** Measured thermosensitivity values (Δ*H)* of TRPV1 temperature studies.

**Supplementary Table 2.** Buffer pH stability as a function of temperature.

## Acknowledgments

We thank Charles R. Sanders for the KCNQ1-VSD plasmid.

## Author contributions

W.D.V.H designed the study. M.K., N.J.S, J.K.H, C.M.M, M.A.C, M.L., and W.D.V.H conducted experiments. M.K., N.J.S, J.K.H, M.L., and W.D.V.H. analyzed the data. M.K., N.J.S, M.L., and W.D.V.H. wrote the manuscript. B.R.C. wrote pulse programs for NMR measurements. All authors read and approved the final manuscript.

## Data and materials availability

All data needed to evaluate the conclusions in the paper are present in the paper and/or the Supplementary Materials. Additional data related to this paper may be requested from the authors. NMR assignment data were deposited in the Biological Magnetic Resonance Bank (BMRB entry 27029).

