## Supporting Information for "Evidence that the TRPV1 S1-S4 Membrane Domain Contributes to Thermosensing"

**Contents**

**Materials and Methods**

*Details of the thermodynamic framework for interpreting temperature-dependent studies*

**Supplementary Figures S1-S7**

*Supplementary Figure S1: The expression and NMR amide backbone assignment of the hV1-S1S4*

*Supplementary Figure S2: Temperature-dependent data of TRPV1 and hV1-S1S4*

*Supplementary Figure S3: Additional data supporting the thermodynamic analyses of the hV1-S1S4 using various biophysical methods*

*Supplementary Figure S4: Supporting data for RDC and PRE NMR measurements*

*Supplementary Figure S5: Mapping of the residues that show temperature-dependent movement*

*Supplementary Figure S6: The hTRPV1-R557A mutant becomes insensitive to both temperature and capsaicin, but the hV1-S1S4-R557A retains the temperature sensitivity*

*Supplementary Figure S7: Structural examples of S4 helix motions in TRP channel activation*

**Supplementary Tables 1 and 2**

*Supplementary Table 1: Measured thermosensitivity values (ΔH) of TRPV1 temperature studies*

*Supplementary Table 2: Buffer pH stability as a function of temperature*

**Materials and Methods**

*Details of the thermodynamic framework for interpreting temperature-dependent studies*

NMR resonance intensities are inversely proportional to the resonance linewidth ($\Delta\upsilon_{FWHH}$, full-width at half-height). For the Lorentzian shaped signals observed in NMR, the intensities and linewidths reflect the transverse relaxation rates ($R_2$) of the molecule, which can be estimated from the linewidth, $\Delta\upsilon_{FWHH} = R_2/\pi$ (*1*). $R_2$ is in turn sensitive to the dynamic properties of the protein in solution, including nanosecond timescale rotational motion, and picosecond to millisecond internal motions, including protein conformational change. Thus, NMR peak intensity should report on the conformational state of a protein. In the hV1-S1S4 temperature-dependent NMR studies we leverage the common framework employed in CD to assess the thermosensitivity in terms of change in enthalpy ($\Delta H$) as described below.

Far-UV CD is commonly used to measure the change in enthalpy ($\Delta H$) between folded or conformational states (*2*). This assumes that the change in measured ellipticity as a function of temperature is directly proportional to the change in protein states. Early studies have shown for two state behavior that there is good correlation between calorimetric and CD measurements of thermodynamic values (c.f.) (*3*). To evaluate the $\Delta H$ for the transition between protein states, one generally assumes a two-state model. In the context of TRP channels this is certainly an oversimplification, but adequately captures accurate thermodynamic values, as has been noted in previous studies (*4*). Briefly, for temperature-dependent equilibrium, CD data at a given wavelength, usually at 222 nm for helical proteins, exhibit clear changes between two states (i.e. the data are sigmoidal in nature). At a given temperature the relative concentrations of conformational states 1 and 2 will be related to the equilibrium constant $K$, which can be written as the fraction of the concentration of conformational state 2 ($\alpha$):

$$K = \frac{[State\ 2]}{[State\ 1]} = \frac{\alpha}{1-\alpha} \qquad [1]$$

$K$ is also related to the change in standard state free energy ($\Delta G°$) between conformational states and noted in the Gibbs-Helmholtz equation:

$$\Delta G° = -RT\ ln\ K \qquad [2]$$

which is in turn is related to the changes in enthalpy ($\Delta H°$) and entropy ($\Delta S°$) between states:

$$\Delta G° = \Delta H° - T\Delta S° \qquad [3]$$

At the midpoint of the transition between temperature-dependent conformational states ($T_{50}$), or the melting temperature ($T_m$) in folding studies, the concentrations of the two structural states are equal and $K = 1$ and $\Delta G° = 0$, resulting in the following identity:

$$\Delta S° = \frac{\Delta H°}{T_{50}} \qquad [4]$$

Combining equations 2, 3, and 4, $K$ can be written in a modified form of the van't Hoff equation using only terms of temperature ($T$), midpoint temperature ($T_{50}$), and the change in enthalpy ($\Delta H°$):

$$K = e^{\left[\left(\frac{\Delta H^{\circ}}{RT}\right)\left(\left(\frac{T}{T_{50}}\right)-1\right)\right]} \quad [5]$$

Rearrangement of Equation 1 gives:

$$\alpha = \frac{K}{(1+K)} \quad [6]$$

A mathematical description of the assumption above, that the observed ellipticity at a given temperature ($\theta_{obs}$) is directly proportional to the ellipticity of conformational states 1 ($\theta_1$) and 2 ($\theta_2$) gives the following relationship:

$$\theta_{obs} = \alpha(\theta_1 - \theta_2) + \theta_1 \quad [7]$$

To calculate the values of $\Delta H$ and $T_{50}$ that best describe the experimental data, initial estimates of $\Delta H$, $T_{50}$, $\theta_1$, and $\theta_2$ are used as initial parameters to non-linear least squares fitting of Equation 5, 6, and 7 to the raw data. The authors used SigmaPlot to perform non-linear least squared fitting of the CD data which resulted in a $\Delta H = 19 \pm 1$ kcal/mol and $T_{50} = 37 \pm 1$ °C.

The method described above assumes that change in the heat capacity ($\Delta C_p$) is negligible. In the context of TRP channel thermosensors, it has been suggested that this may not be the case (*5*). Indeed, Chanda and coworkers have used protein design principles that attempt to increase the magnitude of $\Delta C_p$ for the Shaker voltage-gated potassium channel, a non-thermosensing ion channel, to successfully convert it to a thermosensing ion channel (*6*). Chanda's study clearly indicates that modifying the change in heat capacity can perturb the thermosensitivity of a given protein. However, for wild-type TRP channels at biologically relevant temperatures, it seems that $\Delta C_p$ is relatively small in magnitude. When $\Delta C_p$ changes as a function of temperature then both $\Delta H$ and $\Delta S$ will change significantly as a function of temperature (*5, 7*). However, as reviewed elsewhere (*8*), existing studies of TRP channel thermosensitivity suggest that $\Delta H$ (and thus $\Delta S$) are fairly constant as a function of temperature indicating that $\Delta C_p$ is small in magnitude (*4, 8-12*).

The above described method is completely general for extracting $\Delta H$ from equilibrium data that can be approximated by a simple two state model. To this end, these same procedures were used to estimate the $\Delta H$ from temperature-dependent whole-cell patch-clamp electrophysiology steady state currents (Fig. 2A), where $\theta_1$, and $\theta_2$ were modified to maximal and minimal current values. The resulting electrophysiology values are consistent with previously published TRPV1 thermodynamic values (Table S1). Similarly, NMR resonance intensity as a function of temperature exhibit two-state behavior which was also fit to the above equilibrium thermodynamic model, yielding a $\Delta H$ value that is similar in magnitude to the CD determined value from the hV1-S1S4 (Fig. 2B and fig. S3E). Lastly, this general two-state equation was also used to interpret the temperature-dependence of the intrinsic tryptophan fluorescence in terms of the average emission wavelength (⟨λ⟩), which yielded values of hV1-S1S4 that are consistent with both NMR and CD.

1. J. F. Cavanagh, W. J.; Palmer III, A. G.; Rance, M.; Skelton, N. J., *Protein NMR Spectroscopy: Principles and Practice (Second Edition)*. (Elsevier Academic Press., 2007).

### Supplementary Figures S1-S7

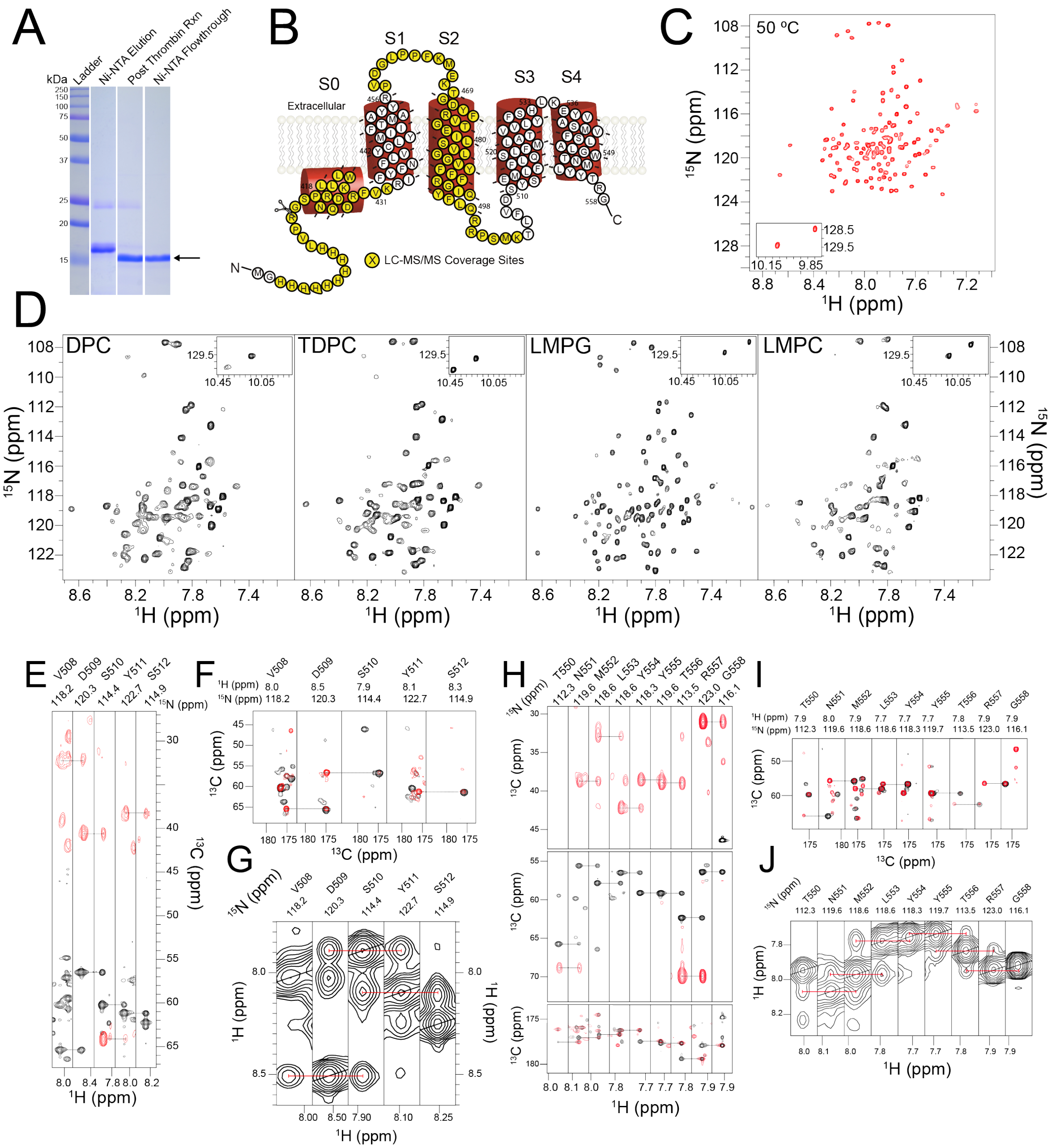


**Supplementary Figure 1. The expression and NMR amide backbone assignment of the hV1-S1S4. (A)** SDS-PAGE gel of the hV1-S1S4 samples of Ni$^{2+}$-NTA affinity chromatography purification, post-thrombin reaction, and 10×His-tag cleaved hV1-S1S4 (lanes 2, 3, and 4 respectively). (**B**) The highlighted residues on the hV1-S1S4 membrane topology represent the coverage for LC-MS/MS as a means to verify the identity of hV1-S1S4 (51% amino acid coverage). (**C**) hV1-S1S4 reconstituted in LPPG micelles results in a high quality $^{1}$H-$^{15}$N TROSY-HSQC NMR spectrum at elevated temperatures (50 °C). (**D**) Optimization of the micellar conditions for the hV1-S1S4 using NMR. To identify a suitable membrane mimic for hV1-S1S4, five membrane mimics were tested. LPPG was chosen to be the most suitable membrane mimic for the hV1-S1S4 as shown in (**C**) after analyzing the number of resonances observed, spectral resolution, and homogeneity of resonance intensities. (**E**) Residues in the TRPV1 S2-S3 loop have been implicated in vanilloid ligand binding and activation. A strip plot of HNCA and HNCOCA from V508 to S512 in the S2-S3 loop is shown on the left (**E**), and this assignment is confirmed by the 4D HNCA and HNCOCA (**F**) and

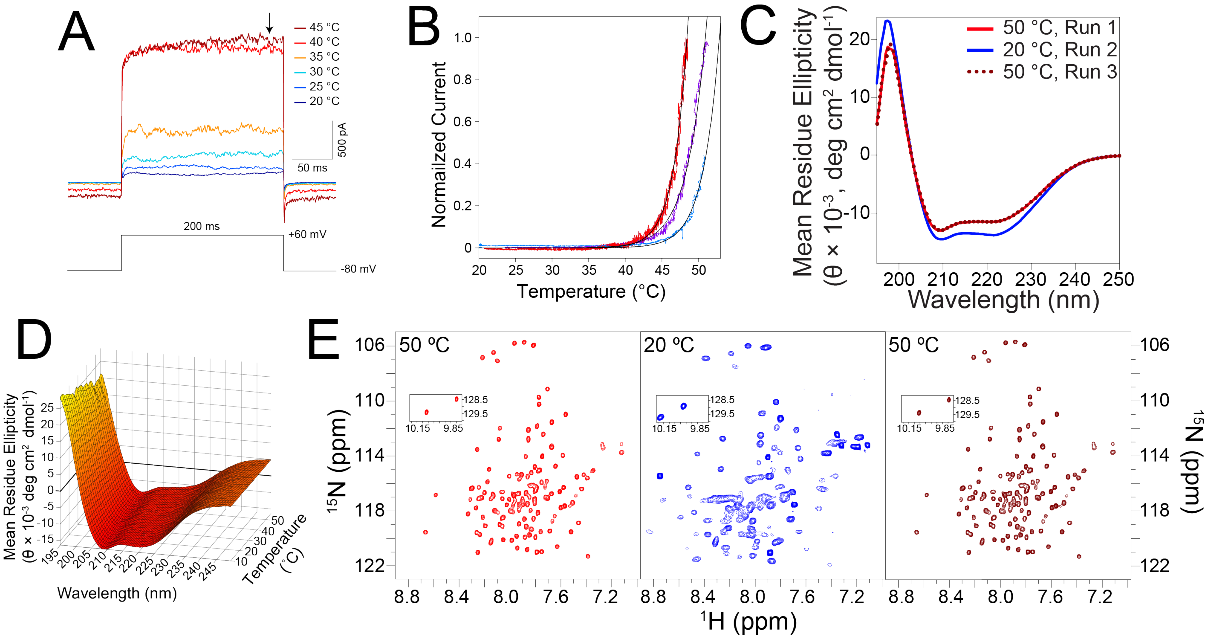


**Supplementary Figure 2. Temperature-dependent data of TRPV1 and hV1-S1S4.** (**A**) The electrophysiology temperature measurement of the human TRPV1. Increased temperature evoked increased current magnitudes, and the data points at the black arrow were plotted against the temperature to generate the plot in Fig. 2A. (**B**) Three individual temperature ramp current values obtained from HEK-293 cells transiently transfected with hTRPV1. Cells were subjected to a temperature ramp from 20 to 50 °C over the course of ca. 35 seconds. The *ΔH* values from each line (red, purple and teal) are obtained from fitting to a pseudo-steady state model (see Methods), resulting in values of 106.8 ± 0.4, 80.1 ± 0.3, and 94.3 ± 0.5 kcal/mol, respectively. (**C**) The reversibility of the hV1-S1S4 was tested with CD between 20 °C and 50 °C. The CD spectra recover to mean residue ellipticity when re-heated. The data shown are of a temperature cycle of elevated, decreased, and returned elevated temperature spectra. (**D**) Superimposed circular dichroism spectra from 10 °C to 57 °C for the hV1-S1S4. The spectra show that key α-helical secondary structure features are generally retained as a function of temperature, which is consistent with the NMR and fluorescence data that the hV1-S1S4 remains seemingly structured over this temperature range. (**E**) The thermal reversibility of the hV1-S1S4 tested with NMR between 20 °C and 50 °C. Like the reversibility shown with CD in (**C**), the hV1-S1S4 that underwent the cycle of heating and cooling retains the initial resonances as shown in the first 50 °C spectrum (red, left), and the second 50 °C spectrum (dark red, right), showing that this domain does not exhibit hysteresis.

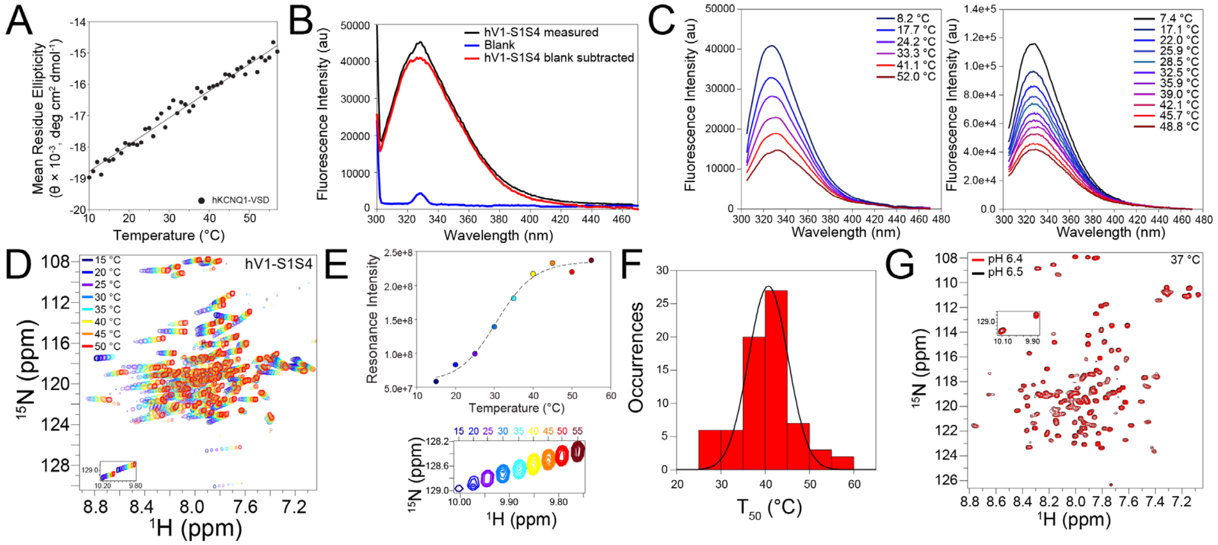


**Supplementary Figure 3. Additional data supporting the thermodynamic analyses of the hV1-S1S4 using various biophysical methods.** (**A**) Non-temperature sensitive human KCNQ1 voltage-sensing domain was used as a negative control. As expected, the human KCNQ1-VSD shows a linear trend in mean residue ellipticity compared to the hV1-S1S4 data, which shows a clear sigmoidal shape. (**B**) The superimposition of the emission spectra of intrinsic tryptophan fluorescence of the hV1-S1S4, buffer (blank) and the hV1-S1S4 black subtracted. (**C**) The superimposition of the emission spectra of the hV1-S1S4 at varying temperature. Two distinct hV1-S1S4 samples were used for these measurements. (**D**) Superimposed $^{1}H$-$^{15}N$ TROSY-HSQC spectra of hV1-S1S4 as a function of temperature show significant chemical shift perturbations. The changes in chemical shift as a function of temperature reflect the thermal expansion of the hydrogen bonding of the secondary structure such that peaks that move less reside in more structured predominantly helical regions. The intensity changes as a function of temperature can be explained by the internal dynamics of the individual amide nuclei, which experience linewidth changes that are suitable for thermodynamic analysis. (**E**) Representative NMR data from W549; as the temperature changes, the intensity data has a sigmoidal shape which reflects a two-state transition and can be fit to extract thermodynamic parameters (see Methods). (**F**) The histogram distribution of melting temperatures ($T_{50}$) calculated from the NMR temperature titration. This was calculated from fitting the sigmoidal curve. The mean $T_{50}$ of 71 resonances was calculated to be 40.7 ± 0.6 °C. (**G**) The $^{1}H$-$^{15}N$ TROSY-HSQC spectra at pH 6.5 and pH 6.4, which is in the range of pH change for the buffer that was used as a function of the temperature ramp, are identical when superimposed. This indicates that temperature induced buffer pH changes are not responsible for the hV1-S1S4 temperature dependent observations noted by NMR and CD experiments.

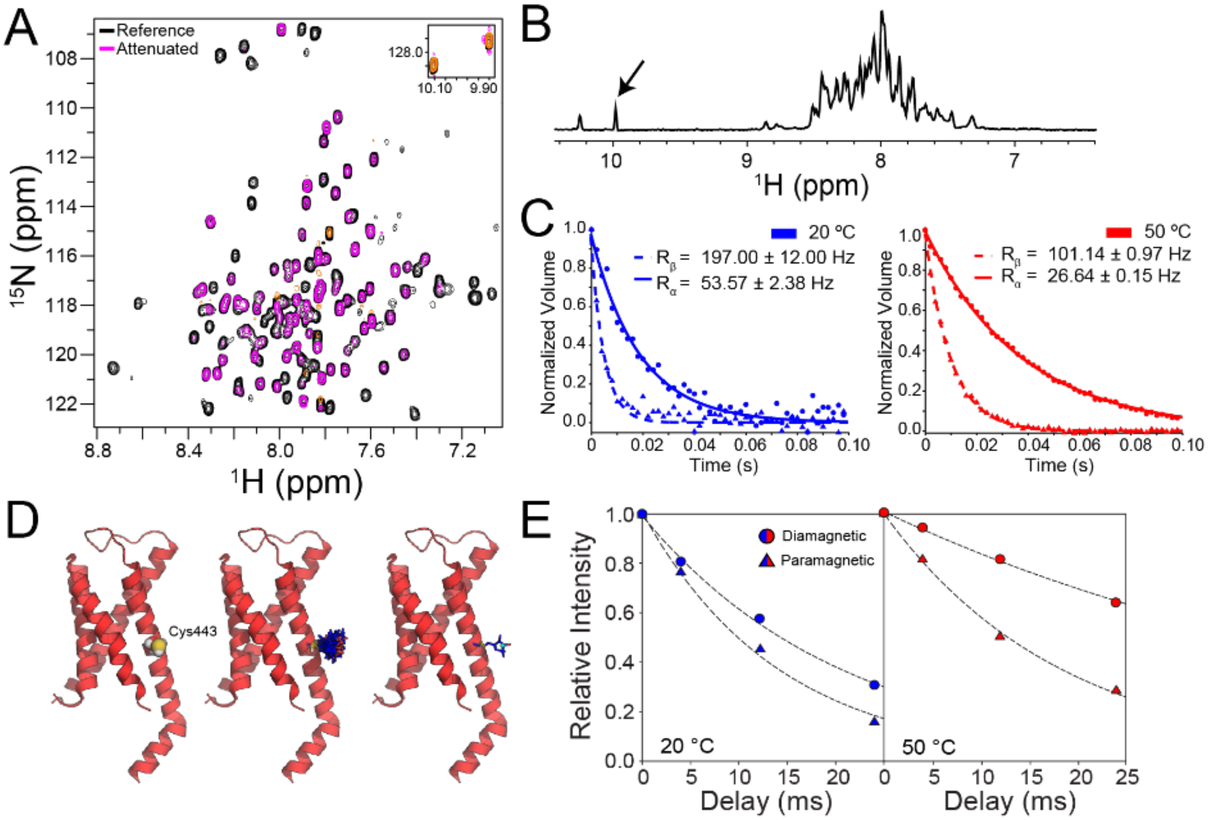


**Supplementary Figure 4. Supporting data for RDC and PRE NMR measurements.** **(A)** The reference (black) and attenuated (magenta) ARTSY spectra of the hV1-S1S4 used for the RDC measurements. (**B**) A representative hV1-S1S4 proton NMR spectrum. The indole amine from the S4 helix residue W549, highlighted with an arrow, was used for analysis in the TRACT experiment (see Methods) due to potential artifacts that can arise from dynamic regions in the protein. (**C**) The relaxation rates for TROSY and Anti-TROSY relaxation states from the W549 indole amine at 20 °C and 50 °C. The relaxation rates were calculated by fitting to a monoexponential decay from which the rates were used to calculate the rotational correlation time ($\tau_c$). (**D**) The location of a lone cysteine in the S1 helix of the hV1-S1S4, C443, was labeled with a maleimide paramagnetic nitroxide spin label, MTSL. The middle structure represents the MTSL-labeled S1-S4 domain (rTRPV1. PDB ID: 5IRZ) computationally, with 85 rotameric states of the MTSL modeled in PyMol (see Methods) and right structure displaying the centroid position of all the rotameric states of the MTSL. (**E**) The representative proton relaxation curves for paramagnetic and diamagnetic states at 20 °C and 50 °C for W427.

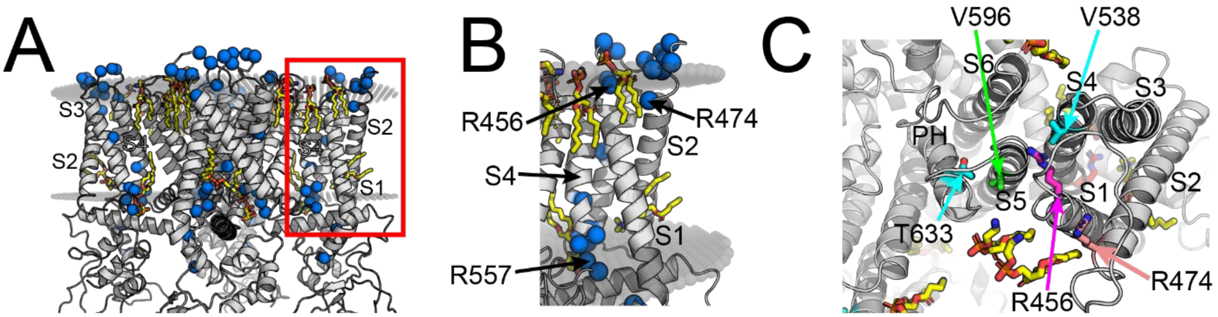


**Supplementary Figure 5. Mapping of the residues that show temperature-dependent movement.** (**A**) The structure of TRPV1 (PDB ID: 5IRZ) in OPM membranes that highlights the residues that show increased solvent accessiblity at 45 °C, highlighted with marine. The main chains of the residues with increased solvation are shown in spheres. The S1-S4 domain is displayed in red box. The lipid molecules are shown in yellow sticks. (**B**) A close-up view of the S1-S4 domain. It is clear that the R456 and R557 are embedded in the membrane. All three residues, R456, R474 and R557, have high values of $\Delta NOE_{45°C-20°C}$ and appear to have temperature-dependent movements. (**C**) Top view of the TRPV1 structure (PDB ID: 5IRZ) that highlights key residues in heat and proton activation. R456 (S1 helix, magenta) is known to be crucial in channel function and gating and is interacting with V538 (S4 helix, cyan). V538 and T633 (pore helix, cyan) are important for proton activation. V596 (S5 helix, green) is a potential key residue for temperature activation, and this residue is surrounded by R456, V538 and T633.

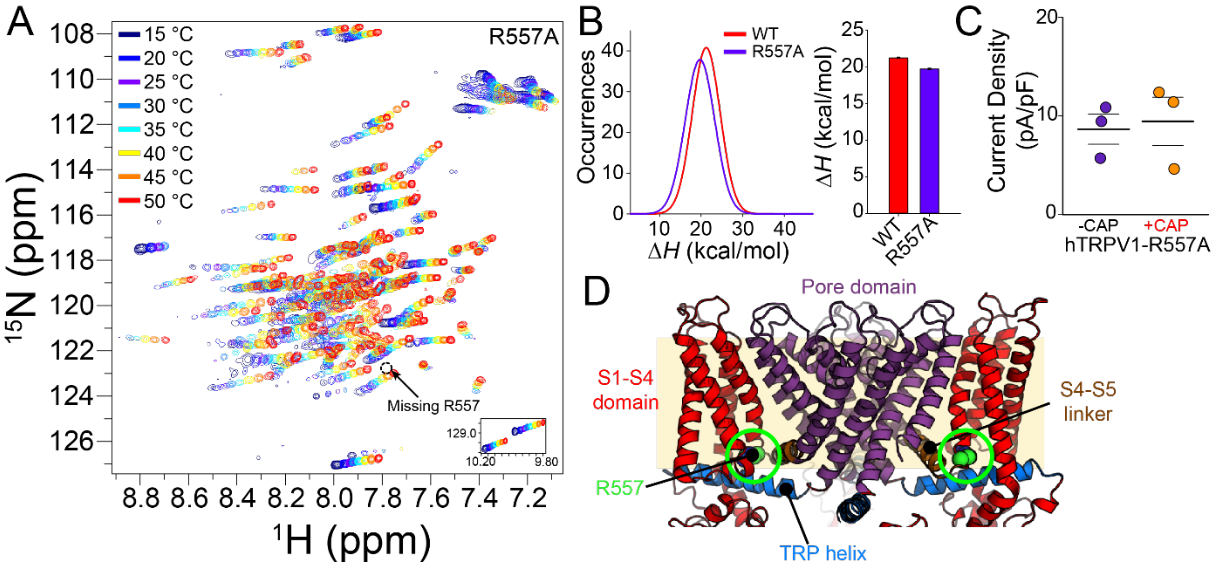


**Supplementary Figure 6. The hTRPV1-R557A mutant becomes insensitive to both temperature and capsaicin, but the hV1-S1S4-R557A retains the temperature sensitivity.** (**A**) Superimposed $^{1}$H-$^{15}$N HSQC spectra of the hV1-S1S4-R557A at temperatures from 15 to 50 °C. As expected the R557 resonance is missing in the R557A mutatnt spectra. (**B**) Comparison in $\Delta H$ between the WT and the hV1-S1S4-R557A mutant. The Gaussian fittings of the $\Delta H$ histograms (left) show a leftward shift, indicating that the R557A (purple line) is slightly less temperature sensitive than the WT (red line), consistent with the loss of a cation-$\pi$ interaction. The ensemble average enthalpy from the respective WT and R557A Gaussian fits and resulting variance are shown in a bar plot (right). (**C**) The jittered plot of hTRPV1-R557A of whole-cell patch-clamp electrophysiology in the presence and absence of capsaicin. Consistent with R557A acting as a coupling mutation, the mutant is insensitive to capsaicin activation at a concentration (1 µM) that would saturate WT hTRPV1. (**D**) Structure of the rat TRPV1 (PDB ID: 5IRZ) that highlights each structural domain. The S1-S4 domains are highlighted in red. The PD that was implanted to the Shaker is demonstrated in purple. We note that R557 (highlighted in green) is not included in the chimeric study, and the phenotypes of this residue indicate that R557 is central to coupling the S1-S4 domain with the PD in channel gating.

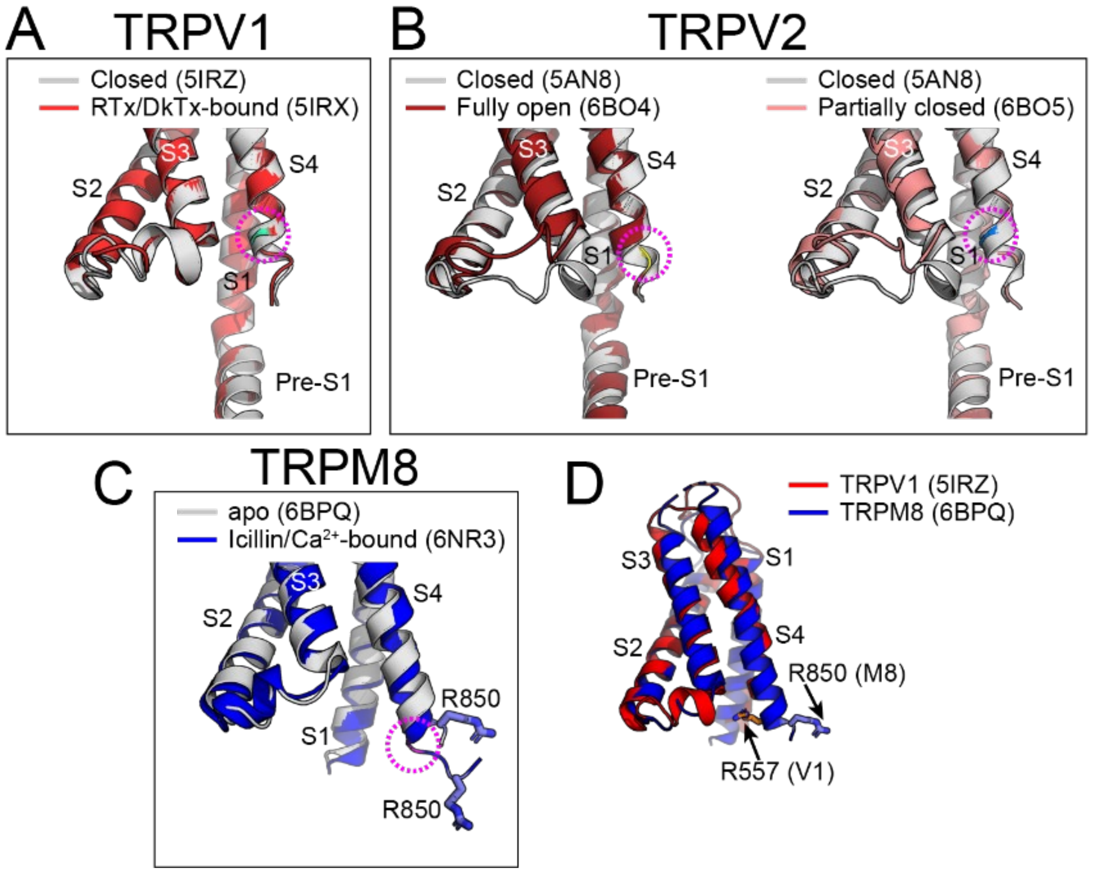


**Supplementary Figure 7. Structural examples of S4 helix motions in TRP channel activation.** (**A**) Superimposed structures of TRPV1 in its resting (PDB ID: 5IRZ, light grey) and active states (RTx/DkTx-bound, PDB ID: 5IRX, red). In the active state, starting around Y555 (light blue, highlighted with magenta circle), the bottom of the S4 helix starts unfurling. (**B**) The same unfurling trend found in TRPV2. The fully-closed structure of TRPV2 (PDB ID: 5AN8) is superimposed with the fully open state that includes the pore turret (PDB ID: 6BO4, dark red on the left), and partially closed state which does not include the pore turret (PDB ID: 6BO5, pink on the right). In both alignments, the unfurling of the S4 begins at the bottom of the S4 helix (highlighted in yellow and marine, with magenta circle). (**C**) Superimposed structures of the apo TRPM8 (PDB ID: 6BPQ, light grey), and the icillin/$Ca^{2+}$-bound TRPM8 (PDB ID: 6NR3, blue). In TRPM8, the bottom of the S4 helix undergoes an extreme conformational change, induced by the $\alpha$- to $3_{10}$- helix transition, starting at the residue highlighted in magenta. This difference can be verified by the displacement of R850, as it travels further down the membrane when TRPM8 is activated. (**D**) The alignment of the S1-S4 domains of TRPV1 (red) and TRPM8 (blue). Although the orientation is different, R557 in TRPV1 and R850 in TRPM8 are located in the C-terminal end of the S4 helix, near the membrane bilayer.

**Supplementary Table 1.** Measured thermosensitivity values (Δ*H)* of TRPV1 temperature studies.

| **Δ*H* (kcal/mol)** | **Potential (mV)** | **Method** | **TRPV1 species** | **Protein Origin/ Membrane Type** | **Reference** |
|---|---|---|---|---|---|
| 98 ± 12 | +60 | Whole-cell | Human | HEK293/HEK293 | **This Study (Steady state)** |
| 94 ± 8 | +60 | Whole-cell | Human | HEK293/HEK293 | **This Study (Temp ramp)** |
| 64.9[a] | -70 | Whole-cell | Rat | HEK293/HEK293 | **Vlachová et al., 2003** |
| 150 ± 13 | -60 | Inside-out | Rat | *X. laevis/ X. laevis* | **Liu et al., 2003** |
| 26.8 | +80 | Inside-out | Murine | HEK293/HEK293 | **Yang et al., 2010** |
| 101 ± 4<br>65 ± 6 | -60, +60 | Outside-out | Rat | HEK293/HEK293 | **Yao et al., 2010** |
| 90 ± 3 | -60 | Whole-cell | Rat | HEK293/HEK293 | **Yao et al., 2011** |
| 86.2 ± 3.9[b] | -60 | Proteoliposome patch | Rat | Sf9 Insect cells/Soybean polar lipids | **Cao et al., 2013** |
| 155 | +100 | Single-channel planar lipid bilayers | Rat | HEK293/3:1 POPC:POPE | **Sun and Zakharian, 2015** |
| 65 ± 5.7[a] | -60 | Whole-cell | Rat | *X. laevis/ X. laevis* | **Zhang et al., 2017** |
| 88 ± 8 | +60 | Inside-out | Rat | HEK293/HEK293 | **Sánchez-Moreno et al., 2018** |

[a] *The reported temperature coefficient ($Q_{10}$) was converted to ΔH according to* $\Delta H \approx 20 ln Q_{10}$ *(5).*
[b] *Protein was purified and reconstituted in proteoliposomes.*

**Supplementary Table 2. Buffer pH stability as a function of temperature.**

| Temperature (°C)[a] | pH[b] |
|---|---|
| 10 | 6.49 |
| 15 | 6.50 |
| 20 | 6.49 |
| 25 | 6.48 |
| 30 | 6.46 |
| 35 | 6.41 |
| 40 | 6.41 |
| 45 | 6.38 |
| 50 | 6.36 |
| 55 | 6.40 |
| 60 | 6.39 |

[a] *Temperature measured with Fluke 52 II thermometer with 80PK-1 beaded type-K thermocouple; according to the instrument specifications, in this temperature range, should produce a maximum temperature error of ±0.4 °C.*
[b] *This data is for the phosphate buffer used in CD, fluorescence and NMR measurements (see Methods).*
